# Spatiotemporal Mapping and Molecular Basis of Whole-brain Circuit Maturation

**DOI:** 10.1101/2024.01.03.572456

**Authors:** Jian Xue, Andrew T. Brawner, Jacqueline R. Thompson, Tushar D. Yelhekar, Kyra T. Newmaster, Qiang Qiu, Yonatan A. Cooper, C. Ron Yu, Yasir H. Ahmed-Braima, Yongsoo Kim, Yingxi Lin

## Abstract

Brain development is highly dynamic and asynchronous, marked by the sequential maturation of functional circuits across the brain. The timing and mechanisms driving circuit maturation remain elusive due to an inability to identify and map maturing neuronal populations. Here we create DevATLAS (Developmental Activation Timing-based Longitudinal Acquisition System) to overcome this obstacle. We develop whole-brain mapping methods to construct the first longitudinal, spatiotemporal map of circuit maturation in early postnatal mouse brains. Moreover, we uncover dramatic impairments within the deep cortical layers in a neurodevelopmental disorders (NDDs) model, demonstrating the utility of this resource to pinpoint when and where circuit maturation is disrupted. Using DevATLAS, we reveal that early experiences accelerate the development of hippocampus-dependent learning by increasing the synaptically mature granule cell population in the dentate gyrus. Finally, DevATLAS enables the discovery of molecular mechanisms driving activity-dependent circuit maturation.

## INTRODUCTION

The early postnatal period encompasses the remarkable transformation of functionally primitive newborns into sophisticated decision-making individuals. This advancement in brain function is enabled by the establishment of efficient neural circuits.^1^ Neural circuits comprise interconnected neurons spanning brain regions and serve as the fundamental units of neuronal processing.^2^ The formation of neural circuits requires preceding processes such as neurogenesis, migration, axon pathfinding, and target innervation. However, the functional ability of neural circuits is conferred during the final stage of neurodevelopment through the activation and remodeling of synapses, here referred to as circuit maturation.^2–7^ Circuit function determines an organism’s behavioral and cognitive capabilities, and many genes associated with neurodevelopmental disorders (NDDs) have been found to be involved circuit maturation.^8–11^

Circuit maturation epitomizes the complexity and asynchrony of brain development.^12–14^ Different brain regions exhibit varying rates and timings of maturation, reflecting the progressive acquisition of skills across development.^15,16^ For example, circuits involved in sensory and motor functions typically mature earlier than those responsible for higher cognitive functions.^15^ Even within a given brain region, neurons do not mature simultaneously.^17–19^ Such complexity and heterogeneity impede our ability to systematically determine the sequence in which circuits mature, preventing us from effectively pinpointing when and where circuit function is disrupted in NDDs.

Circuit maturation is heavily influenced by neuronal activity.^20^ Over time, sensory-driven activity replaces intrinsically generated spontaneous activity to become the dominant driving force for circuit maturation.^18,21–24^ Activity-dependent transcription orchestrates molecular programs to construct and sculpt synaptic connections between neurons. What follows is a surge in the production of both excitatory and inhibitory synapses, closely coordinated by neuronal activity to result in mature circuits with balanced excitation and inhibition (E/I balance).^25^ In fact, a disrupted E/I balance has consistently been observed among NDDs.^26,27^ Since activity-dependent synaptic development is an essential step in circuit maturation, we reasoned that molecular players that are actively involved in this process can be used to monitor the brain-wide sequence of circuit maturation. Npas4, an immediate early gene (IEG), is particularly suited for this purpose due to its robust and selective induction by neuronal activity, involvement in E/I balance through the activity-dependent development of both excitatory and inhibitory synapses, and crucial roles in experience-dependent brain functions like learning and memory.^28–31^

We have developed a Npas4-dependent reporter system that offers a unique capability to permanently and cumulatively labels neurons once they have been activated during the natural course of *in vivo* development. We name this reporter system DevATLAS (Developmental Activation Timing-based Longitudinal Acquisition System). With longitudinal sampling, advanced tissue-clearing techniques, and whole-brain imaging and mapping, we have constructed the first spatiotemporal map of the whole-brain circuit maturation. This resource can be used to pinpoint when and where circuit development deviates from the norm in NDDs, as we demonstrate in an NDD mouse model. To demonstrate the versatility of DevATLAS, we uncover a circuit mechanism by which early experiences accelerate cognitive development. Single-cell RNA sequencing (scRNA-seq) of DevATLAS animals further offers unprecedented opportunities to uncover molecular mechanisms which drive activity-dependent circuit development *in vivo*. Altogether, DevATLAS provides much-needed insights into the intricate processes underlying functional maturation of the brain.

## RESULTS

### DevATLAS enables longitudinal capture of neuronal activation

DevATLAS permanently labels neurons once they are activated and express *Npas4* during development (Figure 1A), which we refer to as neuronal activation hereafter. The Npas4-Cre mouse was created by inserting a P2A-Cre sequence before the stop codon of the *Npas4* coding region. Due to the self-cleavage property of the P2A sequence, transcription from the *Npas4* locus now generates two separate proteins that function independently: full-length Npas4 and Cre. DevATLAS is created when the Npas4-Cre mouse is crossed with either Ai14 or Ai75 mice,^32^ to permanently label neurons with cytosolic (Npas4-Cre;Ai14) or nuclear-localized (Npas4-Cre;Ai75) tdTomato (tdT) following the initial *Npas4* induction during development.

**Figure 1.**
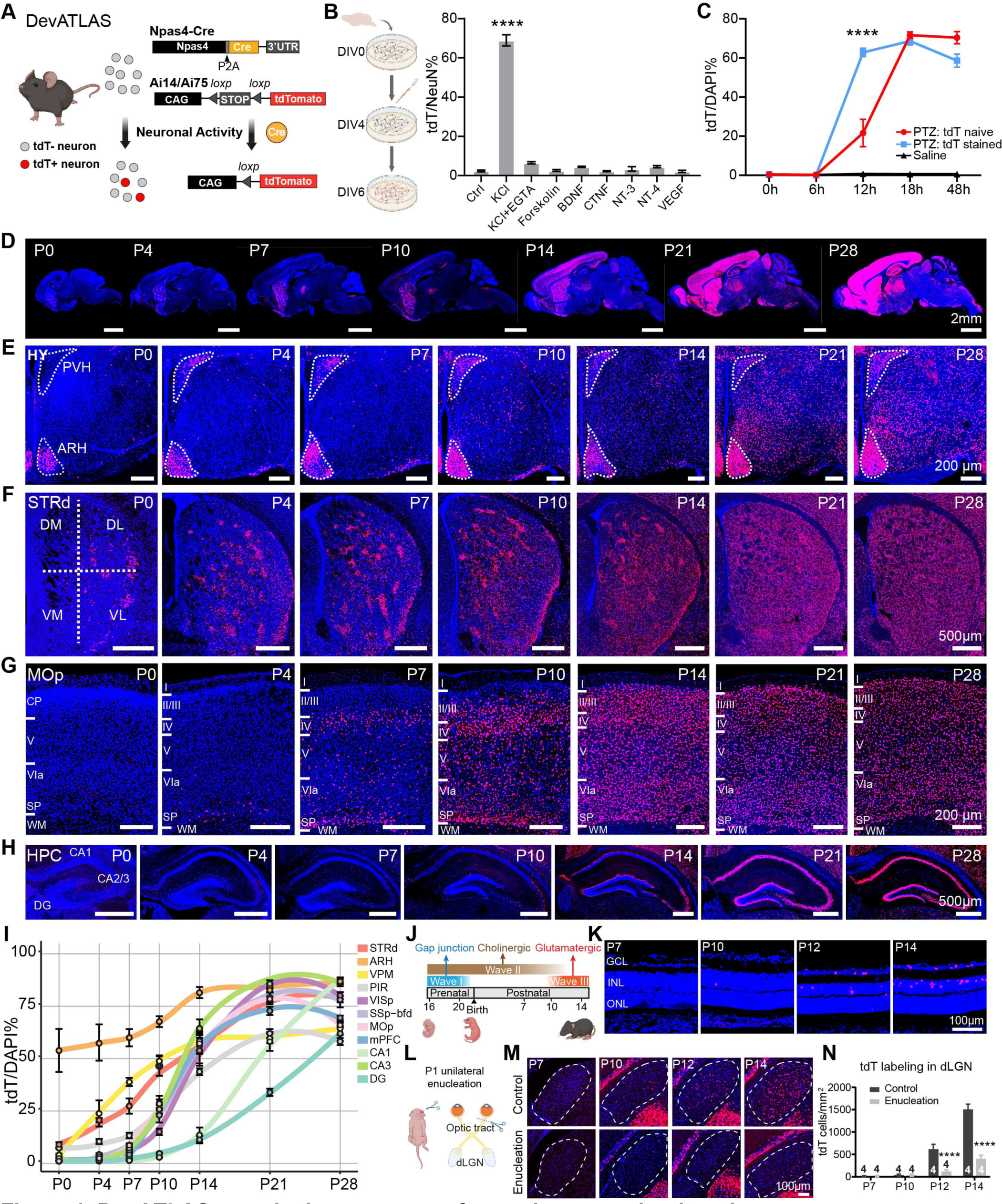
DevATLAS reveals the sequence of asynchronous circuit activation. (A) The design of DevATLAS. (B) DevATLAS (Npas4-Cre;Ai14) signal is selectively triggered by neuronal activity. Cultured hippocampal neurons were treated with extracellular stimuli at DIV4 and the percentage of tdT-labeled neurons (NeuN+) was quantified at DIV 6. tdT, tdTomato. (C) The time course of the appearance of tdT-labeled neurons in CA1 after Pentylenetetrazol (PTZ)-induced seizure (100mg/kg) in P7 DevATLAS (Npas4-Cre;Ai75) mice. (D) Parasagittal brain sections of DevATLAS (Npas4-Cre;Ai75) mice showing tdT labeling at seven developmental ages. (E-H) tdT labeling in hypothalamus (HY) (E), striatum dorsal region (STRd) (F), primary motor area (MOp) (G), and hippocampus (HPC) (H) at seven developmental ages. PVH, paraventricular hypothalamic nucleus; ARH, arcuate hypothalamic nucleus; WM, white matter; SP, subplate; CP, cortical plate; DG, dentate gyrus. Division of the STRd into four quadrants: DL, dorsal lateral; VL, ventral lateral; DM, dorsal medial; VM, ventral medial. (I) Quantification of tdT labeling across 11 brain regions at seven developmental ages. VPM, ventral posteromedial nucleus of the thalamus; PIR, piriform cortex; VISp, primary visual cortex; mPFC, medial prefrontal cortex. n=4-5 mice per age. (J) Schematic depicting the three waves of spontaneous retina activity during development. Modified according to Bansal et al., 2000.^39^ (K) Representative images show tdT labeling in mouse retinas from P7 to P14. GCL, ganglion cell layer; INL, inner nuclear layer; ONL, outer nuclear layer. (L) Diagram for unilateral enucleation, performed at P1. (M and N) Representative images (M) and quantification (N) showing decreased tdT labeling in the contralateral dorsal lateral geniculate nucleus (dLGN) following the enucleation, as compared to the ipsilateral side (Control). Data are shown as mean ± SEM. ****p < 0.0001. See also Figure S1 and statistical data in Table S1.

Like the endogenous *Npas4* gene, DevATLAS tdT labeling is selectively induced by Ca^2+^ influx following neuronal activation and is insensitive to other signaling pathways that are known to activate many other IEGs (Figure 1B).^28,29^ To determine the time course of tdT labeling after neuronal activation, we quantified DevATLAS signal following pentylenetetrazol (PTZ)-induced seizure, to trigger Npas4 expression. In P7 DevATLAS mice, all activated cells were labeled by tdT between 6 and 12 hours after seizure (Figures 1C and S1A). All labeled cells were neurons, a feature of *Npas4* that is not shared by some other IEGs, such as *Fos* (Figures S1B and S1C).

### DevATLAS reveals the brain-wide sequence of neuronal activation

We next monitored DevATLAS tdT labeling to reveal how neuronal activation emerged across the developing brain. DevATLAS revealed the asynchronous onset of neuronal activation across brain regions throughout development (Figure 1D). Regions necessary for survival, such as the arcuate hypothalamic nucleus (ARH, e.g., hunger and satiety) and paraventricular hypothalamic nucleus (PVH, e.g., food intake), were among the first to exhibit robust tdT signal (Figure 1E).^33,34^ Sensory- and motor-associated regions followed, starting with the striatum dorsal region (STRd), which contributes to the integration of sensorimotor response and motor ability (Figure 1F). DevATLAS tdT signal began to emerge in cortical areas about one week later (Figure 1G). Lastly, tdT labeling appeared in brain regions which are dispensable for survival but essential for important higher-level functions (e.g., the hippocampus) (Figure 1H). Such patterns of progressive circuit activation according to changing organismal needs and complexity match the known order of functional development. DevATLAS revealed that areas within ontologically distinct brain regions also exhibit progressive maturation of circuits according to the advancing organismal development. For example, tdT labeling in the hindbrain appeared first in regions critical for survival (parabrachial nucleus, e.g., autonomic functions) (Figure S1D) before progressing to regions regulating sensory and motor function (e.g., pontine gray and superior olivary complex) (Figure S1E).^35^

Quantification of DevATLAS signal in 11 brain regions at seven ages between P0 and P28 substantiated our observations of asynchronous neuronal activation (Figures 1I and S1F). At this more detailed resolution, differences in cortical maturation sequence are apparent. For example, the olfactory system develops earliest among sensory systems and accordingly, the piriform cortex (PIR) exhibited increased tdT labeling at P0 and P4 compared to other primary sensory cortices (Figure S1F). The slight delay in maturation of the visual cortex (VISp) compared to other primary sensory cortices is also captured by DevATLAS (Figure S1F). Interestingly, the initial neuronal activation of the medial prefrontal cortex (mPFC) and the CA3 region of the hippocampus, both of which regulate higher-order behaviors that emerge later in young adulthood, coincided with activation of the primary sensory and motor cortices and plateaued a week or more before other higher-level brain regions in the hippocampus (e.g., the CA1 and dentate gyrus) (Figure S1F). This observation suggests that the behavioral and cognitive complexities attributed to the mPFC and CA3 are not limited by the neuronal activation of those regions, per se.

Altogether, DevATLAS reveals that brain regions are sequentially activated in a general trend according to physiological needs: those crucial for basic survival emerge first, followed by regions which interface with the external environment (motor and sensory), and finally regions which integrate and interpret external environmental signals for higher-order behavioral execution. The sequence of tdT labeling coincides with the order of functional circuit development but implies subtle differences from behavioral maturity in some cases.

As we established in the introduction, circuit maturation is not a singular moment or event but rather a development phase following circuit assembly wherein the connections between neurons are activated and then reinforced. We propose that the neuronal activation captured by DevATLAS is a brain-wide benchmark of the initial stage of circuit maturation that confers advanced synaptic properties and enables behavioral maturity. We next conducted a series of experiments to test this hypothesis, investigating whether: (1) DevATLAS captures synaptic activity *in vivo*; (2) DevATLAS tdT labeling coincides with the timing of circuit assembly; and (3) DevATLAS identifies neuronal populations with advanced functional maturity.

### Synaptic activity triggers DevATLAS labeling during circuit maturation

To confirm that DevATLAS-labeled neurons have experienced synaptic activity *in vivo*, we examined when, where, and which types of neuronal activity triggers tdT labeling in the sensory systems.

The development of the visual system is known to be driven by both spontaneous and, later, light-driven retinal activity that is transmitted to the brain via retinal ganglion cells (RGCs).^22,36–38^ In the retina, tdT labeling emerged during the window of the glutamatergic wave of spontaneous retinal activity around P10, but not the earlier gap junction- and cholinergic-mediated waves (Figure 1J and 1K).^39^ This suggests that Npas4 expression is specifically triggered by glutamatergic synaptic activity. Dark rearing, without any light exposure (Figure S1G), significantly reduced, but did not eliminate, the number of tdT-labeled RGCs by P16 (Figures S1H and S1I). These observations demonstrate that DevATLAS captures both spontaneous and sensory-driven neuronal activity in the retina.

To investigate the effect of RGC activity on the downstream dorsal lateral geniculate nucleus of the thalamus (dLGN), we performed unilateral enucleation to eliminate both spontaneous and light-evoked retinal input from one eye while preserving local connections within the dLGN (Figure 1L). This led to a profound absence of tdT labeling in the contralateral dLGN at P12 and P14 (Figures 1M and 1N). These findings indicate that the activity transmitted from the retina, rather than local dLGN activity, is the main driver of DevATLAS signal within the dLGN. Interestingly, dark rearing, which preserves only spontaneous RGC activity, led to a small but significant decrease in tdT labeling in the dLGN by P16 (Figures S1J and S1K). These results suggest that tdT labeling reflects the crucial role of the spontaneous, glutamatergic retinal wave in shaping synaptic connections between RGCs and dLGN neurons.^22^

In the olfactory system, which develops early and is functional at birth, DevATLAS exhibited labeling in areas associated with olfactory processing by P0, such as the PIR and the main olfactory bulb (MOB) (Figure S1L). Consistent with previous findings, removal of olfactory input with naris occlusion at P0 significantly reduced the number of tdT-labeled neurons in the MOB and PIR at P4 (Figures S1M-S1Q).^40^ This result supports the critical role of spontaneous activity in the development of olfactory circuits.^41^

Taken together, our data demonstrate that DevATLAS captures both spontaneous and sensory-driven glutamatergic activity propagated throughout sensory systems, supporting our hypothesis that DevATLAS captures the activity-dependent initiation of circuit maturation.

### DevATLAS catches patterns of circuit assembly

To establish whether neuronal activation documented by DevATLAS is involved in the wiring of functional circuits, we examined the appearance of tdT labeling in three developmental systems where circuit assembly is well-documented.

The STRd is one of the earliest regions showing robust tdT labeling (Figures 1D and 1F). We found that striosome neurons, known as the early-born striatal population,^42^ were labeled and thus activated earlier (P0-P4) than the later-born matrix neurons (P7-P21) (Figures 1F,2A, and S2A). To investigate whether the early tdT labeling of striosome neurons indicates of their integration into the circuits first,^43,44^ we performed immunohistochemistry experiments at P0. We examined the distribution of axonal terminals from the substantia nigra pars compacta (TH+), the cortex (vglut1+), and the thalamus (vglut2+). Remarkably, axonal terminals from all three regions converged onto the tdT-labeled striosome region and were noticeably absent from the matrix region, which was not labeled by tdT at this time (Figure 2B). At P7, consistent with tdT labeling pattern, all terminals appeared in matrix (Figure S2B). Interestingly, we also observed that the lateral STRd, which receives inputs from the sensorimotor cortex, displayed tdT labeling before the medial STRd, which receives input from higher-order association areas (Figures 1F and 2A).^45^ This progression of tdT labeling likely reflects the order and timing of corticostriatal circuits assembly, which had not been previously demonstrated.

**Figure 2.**
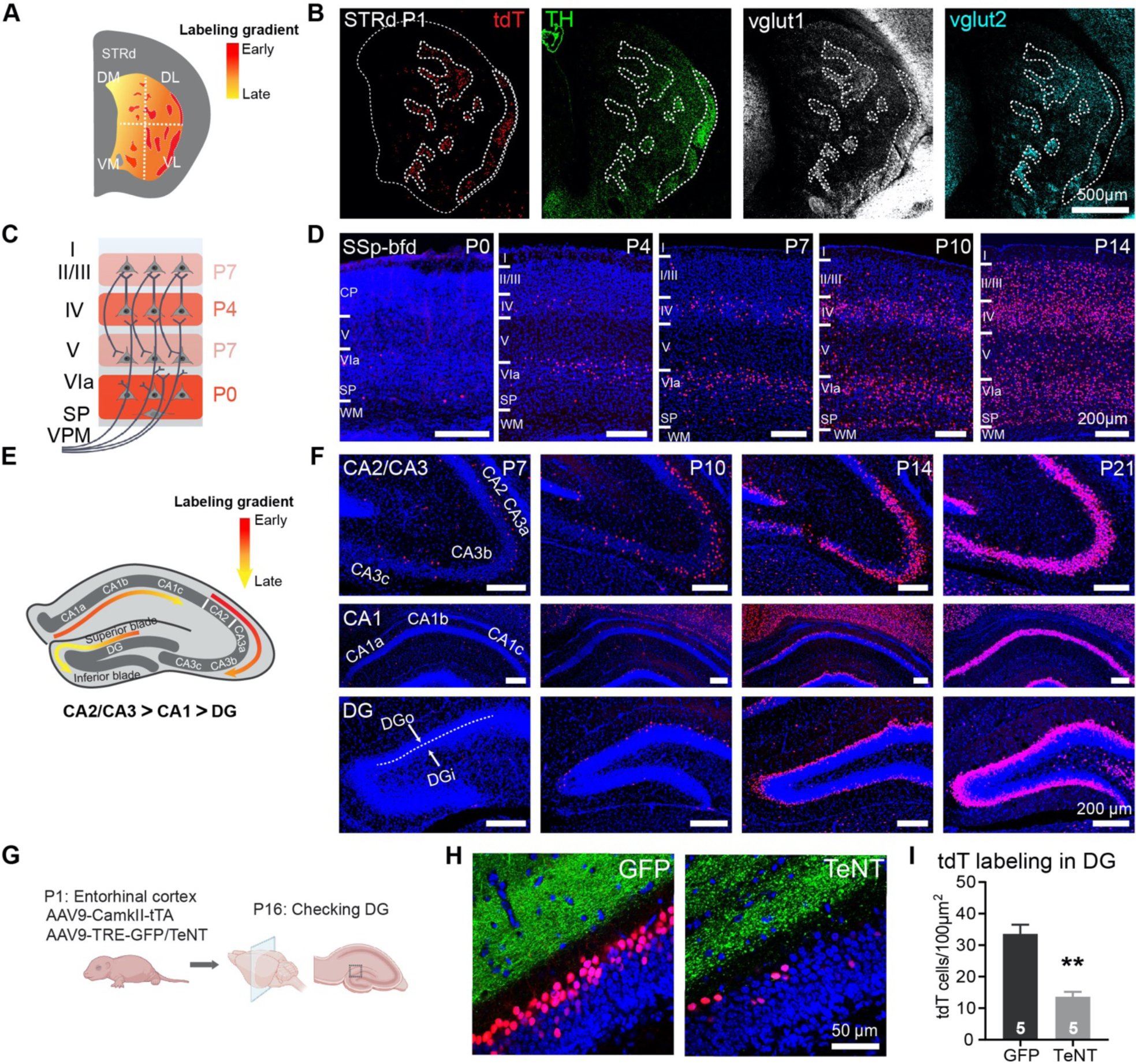
DevATLAS labeling follows the sequence of circuit assembly. (A) Schematic of tdT labeling sequence in STRd. (B) Presynaptic markers co-localize to tdT-labeled striatal regions at P0. Tyrosine hydroxylase (TH) labels the projection terminals of dopaminergic neurons; vglut1 and vglut2 label the projection terminals of glutamatergic neurons in cortex and thalamus, respectively. (C-D) Schematic (C) and representative images (D) of thalamocortical inputs from VPM to primary somatosensory area, barrel field (SSp-bfd) layers during early postnatal ages. The colors indicate the timing of tdT labeling in development with the darker red meaning earlier labeling, and lighter red meaning later labeling. The age of initial tdT labeling is to the right of each layer. (E-F) Schematic (E) and representative images (F) of hippocampal tdT labeling sequence during development. DGo, dentate gyrus outer shell; DGi, dentate gyrus inner core. (G) Experimental scheme. AAVs were injected into entorhinal cortex (EC) of DevATLAS (Npas4-Cre;Ai75) mice at P1 to express tetanus toxin (TeNT) and block neurotransmission from EC to DG. (H and I) Representative images (H) and quantification (I) showing a dramatic reduction in tdT labeling by TeNT, compared to GFP-expressing controls. Data are shown as mean ± SEM. **p < 0.01. See also Figure S2 and statistical data in Table S1.

In the barrel cortex (SSp-bfd), the timing and sequence of DevATLAS signal closely followed the formation of the thalamocortical circuit (Figure 2C). tdT labeling first emerged in SSp-bfd layer VI and then in layer IV, coinciding with the timing when thalamocortical axons heavily innervate these regions (P4-P10) (Figures 2D, S2C, and S2D). Sparse tdT labeling then appeared in layer II/III and layer V at P7-P10, in line with the initial activation of barrel minicolumn circuits (Figures 2C and 2D).^24,46^ Of note, layer V excitatory neurons are born earlier than those in layers IV and II/III yet are labeled later (Figures S2C and S2D), suggesting that the order in which neurons are assembled into circuits and activated does not necessarily correspond to neuronal birth order.

The hippocampus is another region where we observe agreement between the sequence of circuit assembly and DevATLAS signal. tdT labeling first emerged in the CA2/CA3, followed by the CA1 and then the dentate gyrus (DG) (Figures 2E, 2F, and S2E). This pattern recapitulates the sequence of entorhinal cortex-driven hippocampal maturation reported in previous studies.^47^ tdT labeling also reflects the order by which portions of each subregion integrated into the hippocampus circuit: from CA1a/b to CA1c, from CA3a/b to CA3c, and starting from the edge where early-born neurons reside in all hippocampal subregions (Figure 2F).^18^

The patterns of tdT labeling across the developing brain indicate that the sequence of circuit maturation is mainly determined by circuit assembly order and is partially associated with neuronal birth order (Figures 2B, 2D, 2F, and S1F). We utilized DevATLAS to confirm that the propagation of neuronal activity across assembled circuits occurred through synapse. We found that tdT signal in the DG was drastically reduced when presynaptic communication from its major glutamatergic excitatory input, the entorhinal cortex, was ablated by tetanus toxin (Figures 2G-2I). Taken together, DevATLAS signal can be used to capture the sequence of circuit assembly during brain development.

### DevATLAS captures the population that is functionally more mature

If DevATLAS captures circuit maturation in response to the synaptic activation of assembled circuits, we would anticipate that, at the cellular level, the tdT-labeled neurons are more mature and possess stronger synaptic connections than their unlabeled neighboring neurons. To assess this hypothesis across brain regions and cell types, we focused on the STRd, which develops early and exhibits the drastic heterogeneity in tdT labeling shortly after birth (Figure 1F), and the DG, which exhibits tdT labeling weeks later (Figure 1H) and is associated with cognitive function.

In the P0 STRd, we found that approximately 90% of tdT-labeled medium spiny neurons (MSNs), the predominant striatal cell type, expressed DARPP32, a marker of mature MSNs (Figures 3A and 3B).^48^ In contrast, less than 10% of unlabeled MSNs expressed DARPP32. Like most neurons, the resting membrane potential of MSNs decreases as part of neuronal maturation. We found that tdT-labeled MSNs displayed significantly more hyperpolarized resting membrane potential compared to unlabeled MSNs (Figure 3C). In addition, the reversal potential of GABA was significantly lower in tdT-labeled MSNs due to a much lower intracellular chloride concentration, resulting in a more hyperpolarized GABA driving force (Figure 3C). Thus, the developmental GABA switch has occurred in tdT-labeled MSNs but not yet in their unlabeled counterparts. This developmental advancement is key for functional signaling and the integration into mature circuits.^49^ To further examine the maturity, we measured synaptic properties of tdT-labeled and unlabeled MSNs in P7 DevATLAS mice. The tdT-labeled MSNs exhibited significantly higher frequencies of spontaneous miniature excitatory (mEPSC) and inhibitory postsynaptic current (mIPSC) (Figures 3D and 3E). These results strongly indicate that the activation evidenced by DevATLAS signal reflects cellular and synaptic maturity in the STRd.

**Figure 3.**
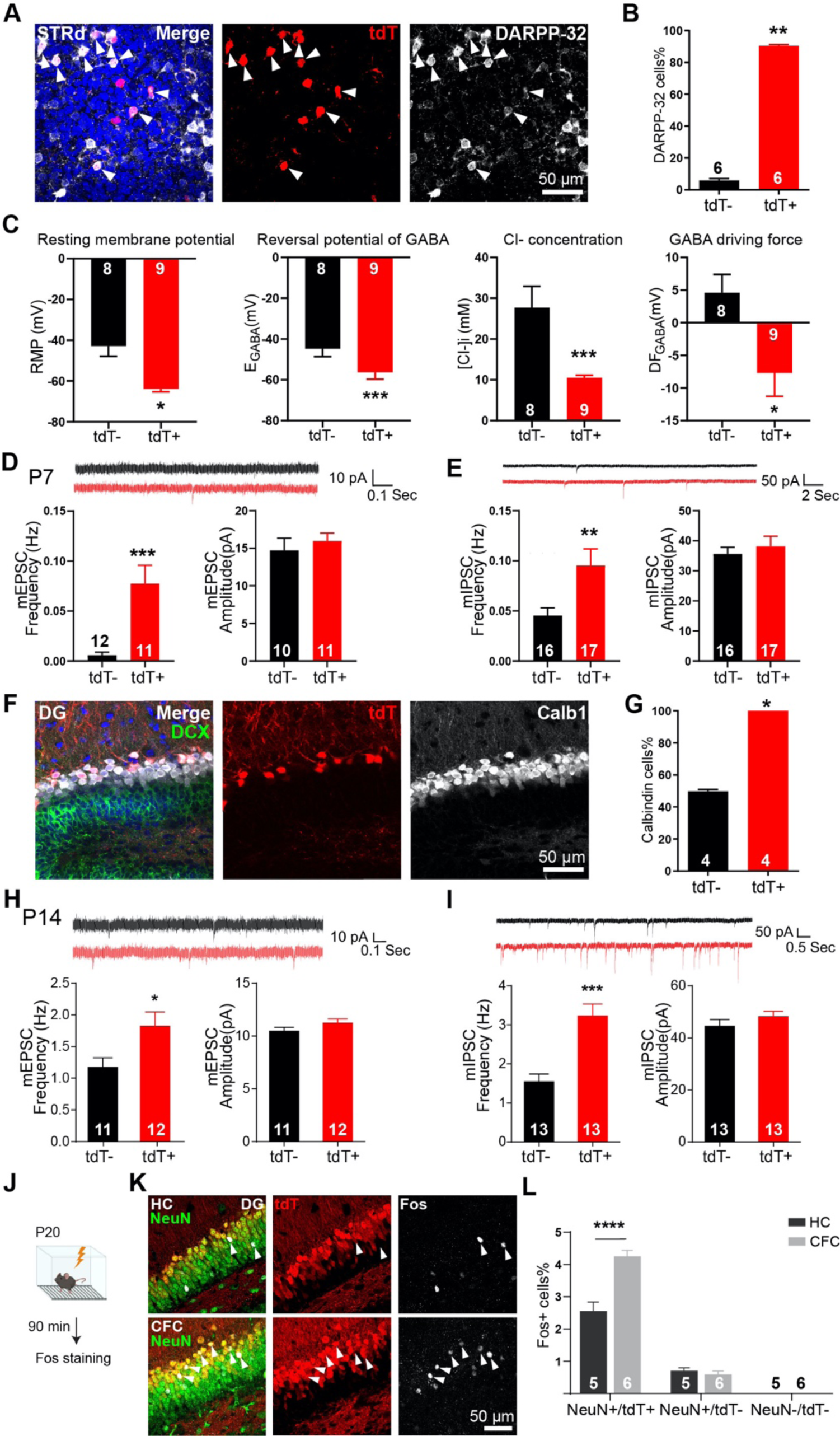
DevATLAS-labeled neurons are functionally more mature. (A and B) Representative images (A) and quantification (B) of the enrichment of mature neurons in tdT-labeled (tdT+) population in the STRd (P0-1) of DevATLAS (Npas4-Cre;Ai14) mice. DARPP-32 is a maturation marker of medium spiny neurons (MSNs). (C) tdT+ MSNs display a significantly more hyperpolarized resting membrane potential, lower reversal potential of GABA responses (E_GABA_), lower cellular Cl^-^ concentration, and higher GABA driving force than tdT-MSNs at P7. (D and E) At P7, tdT+ MSNs showed significantly higher mEPSC (D) and mIPSC (E) frequencies compared to tdT-MSNs. Representative traces are shown on the top. (F and G) Representative images (F) and quantification (G) of the enrichment of mature neurons in tdT+ population in the DG of P14 DevATLAS (Npas4-Cre;Ai14) mice. All tdT+ neurons are expressing granule cell (GC) maturation marker, Calb1 (Calbindin 1). DCX, Doublecortin. (H and I) At P14, tdT+ GCs showed significantly higher mEPSC (D) and mIPSC (E) frequencies compared to tdT-GCs. Representative traces are shown on the top. (J) Schematic of Fos labelling after contextual fear conditioning in the DG of DevATLAS (Npas4-Cre;Ai14) mice. (K and L) Representative images (K) and quantification (L) showing that contextual learning exclusively activates tdT+ neurons at P20. White arrows indicate Fos+/tdT+ neurons. Data are shown as mean ± SEM. *p < 0.05, **p < 0.01, ***p < 0.001, ****p < 0.0001. See statistical data in Table S1.

Similar results were observed in the granule cells (GCs) of the DG. At P14, all tdT-labeled cells in the DG were identified as mature GCs, while less than half of unlabeled GCs expressed the maturation marker Calbindin (Figures 3F and 3G).^50^ Both mEPSC and mIPSC frequencies were significantly higher in tdT-labeled GCs than unlabeled GCs (Figures 3H and 3I). We next examined whether the more abundant synaptic inputs received by the tdT-labeled GCs were related to hippocampal learning and memory ability. We trained P20 DevATLAS mice with contextual fear conditioning (CFC). Neurons acutely activated during CFC were identified by their expression of Fos, a proxy for memory engram neurons (Figure 3J). We found that CFC increased acute neuronal activation exclusively in the tdT-labeled GCs (Figures 3K and 3L), strongly indicating that tdT-labeled neurons are more likely than their unlabeled counterparts to participate in memory encoding.

Taken together, our results demonstrate that tdT-labeled neurons are more mature at the cellular, physiological, synaptic, and behavioral levels. DevATLAS has proven uniquely capable of tagging neurons in different brain regions and circuits that have initiated the circuit maturation in response to synaptic activity. Critically, DevATLAS accomplishes this amidst the asynchronous and heterogeneous *in vivo* environment of the developing brain.

### A spatiotemporal map of circuit maturation across the whole brain

Having established that DevATLAS can capture sequences of circuit maturation in different brain systems and ages, we next sought to create, for the first time, a longitudinal, spatiotemporal map of circuit maturation across the whole brain. Such a resource would allow researchers to identify when and where neuronal activation begins in their region of interest, enabling in-depth analysis of the specific developmental processes. We built a whole-brain imaging and data analysis pipeline to create this much-needed resource for the neuroscience research community (Figure 4A).

**Figure 4.**
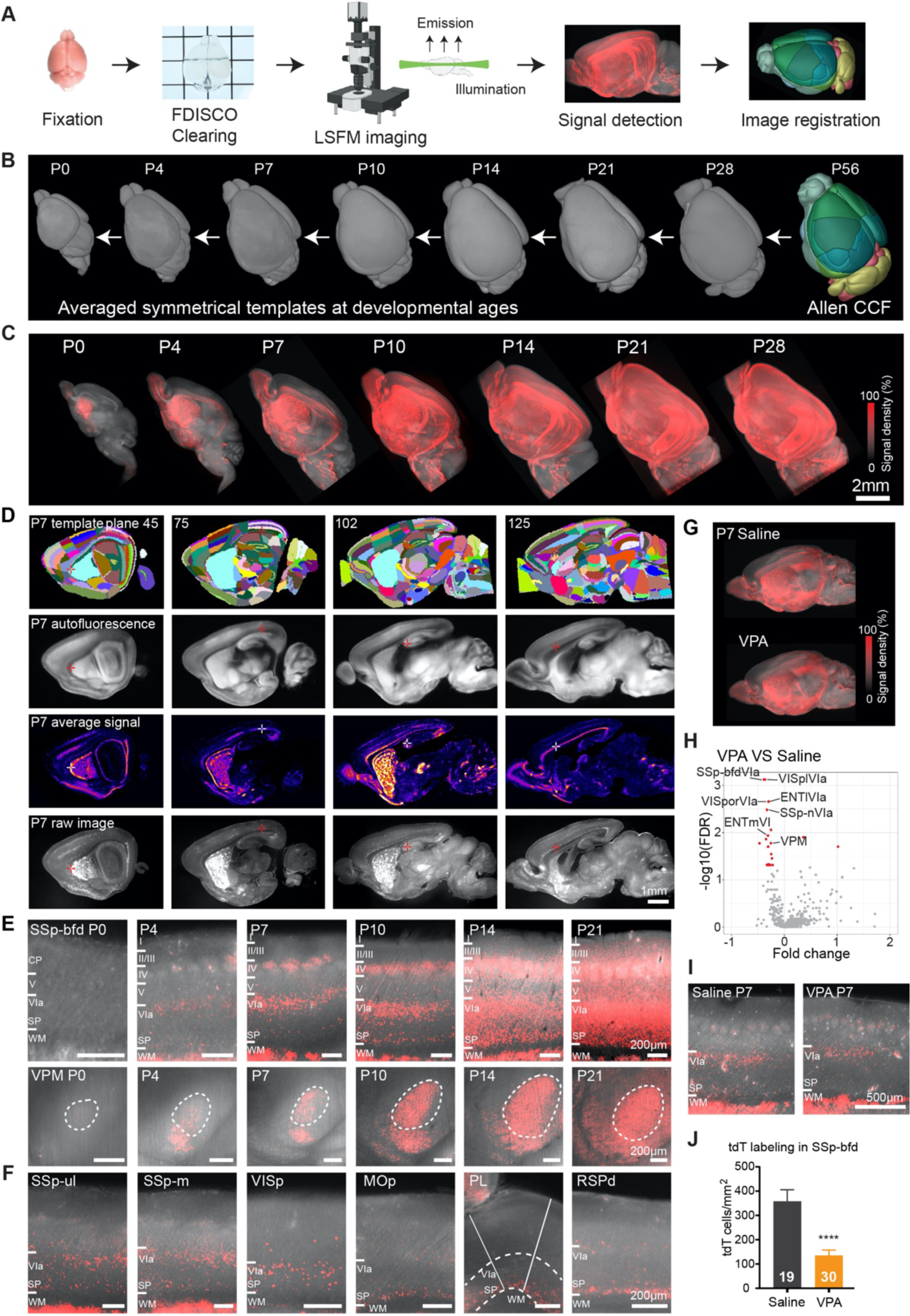
Spatiotemporal mapping of circuit maturation during early development across the whole-brain. (A) Schematic of the whole-brain clearing, imaging, and data analysis pipeline. (B) Developmental brain templates were sequentially registered to establish anatomical labels at each age. (C) Averaged tdT signals overlaid onto age matched template brains. (D) Dataset navigation in the synchronized template registration, autofluorescence, average tdT signal, and raw tdT signal image. The pointers demonstrate the regions of interest across the synchronized images. (E) Representative lightsheet fluorescence microscope (LSFM) images with detected tdT signals in the SSp-bfd and the VPM from P0 to P14. (F) Representative LSFM images showing the tdT labeling different patterns observed in layer VI at P7. SSp-ul, primary somatosensory area, upper limb; SSp-m, primary somatosensory area, mouth; PL, prelimbic area; RSPd, retrosplenial area, dorsal part. (G) Averaged tdT signals in the brains of P7 mice treated with saline or valproic acid (VPA). (H) Volcano plot showing brain regions with different signal density between the saline- and VPA-treated mice. Red dots indicate regions with significant difference (FDR<0.05). n= 19 (saline) and 30 (VPA). (I and J) Representative LSFM images with tdT signal (I) and manual quantification (J) showing VPA treatment decreased tdT labeling in the layer VIa of SSp-bfd. Data are shown as mean ± SEM. ****p < 0.0001. See also Figure S3 and statistical data in Table S1.

First, a series of reference brain templates were generated by adopting a previously established strategy,^51^ which was then adjusted based on our current clearing and imaging method to cover neonatal ages and provide more detailed annotations based on Allen common coordinate framework labels (Figure 4B).^52^ Brain samples were collected from DevATLAS mice at seven developmental time points spanning P0 to P28. Brains were tissue-cleared using FDISCO,^53^ imaged on a LaVision Ultramicroscope II light sheet fluorescence microscope (LSFM), and mapped onto our age-matched reference templates (Figure 4C). Our dataset allows for visualizing the spatiotemporal order of circuit maturation along the sagittal, coronal, and horizontal axes. This visualization allows for simultaneous, coordinate-linked examination of four data channels (Figures 4D, S3A, and S3B, Methods): annotation template, the averaged tdT signal from at least 12 brains collected for each time point, the autofluorescence image (giving a rough anatomical outline of the brain), and an individual sample brain displaying raw tdT signals (Figure 4D).

We highlight the thalamocortical barrel circuit to exemplify how the DevATLAS whole-brain imaging dataset reveals the sequence of circuit maturation. The formation of this circuit, previously described using traditional histology methods (Figure 2D), is plainly evident in LSFM images of the SSp-bfd region from P0 to P21 (Figure 4E). The barrel formations can even be seen within individual brain samples (Figure 4E, P7). Concurrently, LSFM images of the VPM, the thalamic input to SSp-bfd, exhibit tdT labeling closely correlated to that in the SSp-bfd (Figure 4E), consistent with the known role of VPM neurotransmission driving the formation of the thalamocortical barrel circuit.^54^

To gain a more general insight into whole-brain circuit activation patterns, tdT signals were mapped to anatomical brain regions (Table S2) and their volumetric proportions were extracted and then hierarchically clustered across ages and regions (Figure S3C, Table S3). All the raw and processed data, along with the new developmental brain templates, can be freely accessible through Mendeley data and BrainImageLibrary.

One of our goals for the DevATLAS spatiotemporal map is to reveal potentially unknown roles of regions in brain development. Two such regions, the fasciola cincerea and induseum griseum, stood out in the tdT signal clustering data. They are little-studied regions of the hippocampus that exhibited robust tdT labeling by P4, much earlier in development than the remainder of the hippocampus regions, suggesting that these regions may play not-yet-identified roles in circuit maturation during the early developmental stages (Figure S3C). Of note, neuronal activity in the induseum griseum has been shown to be impacted by maternal stimulant abuse during pregnancy, highlighting the potential significance of the activity-dependent development of this region and the need for further investigation.^55^

Another intriguing observation apparent in DevATLAS LSFM images is the distinct patterns of tdT labeling in cortical layer VI across different cortical areas at the same developmental age (Figures 4F and S3D). In the primary sensory cortices such as the somatosensory cortex (SSp) and the VISp, tdT labeling at P7 first appears strongly in upper layer VI (VIa), more so than the lower layer VI (VIb, subplate). In contrast, non-sensory and association cortices such as the primary motor cortex (Mop), prelimbic cortex (PL), and retrosplenial cortex (RSPd), exhibited initial tdT labeling in layer VIb/subplate (Figures 4F and S3D). As layer VI is the first cortical layer to receive thalamus input, these results suggest differences in thalamocortical circuit assembly or maturation across different cortical areas.

Altogether, whole-brain imaging and analysis of DevATLAS provides high-throughput and quantitative measurements of the sequential circuit maturation on the whole-brain scale. The spatiotemporal whole-brain map of neuronal activation offers an unprecedented opportunity for the neuroscience community to trace and explore the steps of circuit maturation during early brain development.

### Whole-brain mapping reveals disruptions in circuit development in a mouse model of autism spectrum disorder

Circuit development is often disrupted in NDDs, such as autism spectrum disorder (ASD).^56^ Identifying where and when in development these perturbations occur has been a bottleneck in NDD research that limits investigation to a few pre-selected brain regions without knowing where and when the synapse-level perturbations are concentrated. To overcome this limitation, we sought to leverage DevATLAS and the high-throughput whole-brain imaging and analysis pipeline as a discovery platform that can systematically identify when and where circuit development is disrupted in NDDs.

To this end, we employed an established NDD model with ASD like behavioral deficit: prenatal valproic acid (VPA) exposure.^57^ Pregnant DevATLAS dams were injected with either VPA or saline at E12.5 (Figure S3E, see Methods). Consistent with previous reports,^58^ VPA-exposed pups exhibited developmental phenotypes consistent with ASD, including impaired physical development and altered social communication at the early postnatal stage (Figure S3F). Whole-brain imaging mapping was carried out using FDISCO-processed P7 brains from both VPA and control saline treatment. Consistent with a lack of gross morphological changes in NDDs, overall tdT labeling appeared similar at P7 (Figure 4G). However, several brain regions exhibited significant changes in tdT densities between saline- and VPA-treated brains, with VPA exposure predominantly reducing tdT signal (Figure 4H, Table S4). Notably, layer VIa of sensory cortices, particularly the SSp-bfd, exhibited the greatest reduction of tdT labeling in VPA-exposed animals (Figures 4H and 4I). These results were confirmed with manual counting of LSFM images (Figure 4J). Interestingly, the VPM also displayed decreased tdT densities (Figure 4H). These data suggest that sensory processing abnormalities in persons with NDD may be linked to early-life changes in sensory circuit maturation.^59^

Thus, DevATLAS, combined with the whole-brain imaging and analysis pipeline, enables the identification of the brain regions and neural circuit maturation where disrupted development may underlie behavioral impairments.

### DevATLAS uncovers experience-driven facilitation of cognitive development

Given DevATLAS’s unique ability to track the neuronal activation history of neurons undergoing activity-dependent functional maturation, we sought to leverage the reporter system to explore the mechanisms by which sensory experience influences circuit maturation in regions like the hippocampus that receive highly processed sensory information. Unlike the sensory systems, experience-dependent maturation of non-sensory brain regions remains under explored.

To manipulate sensory experience, we implemented early environmental enrichment (EE, see Methods) in experimental mice (EE-reared mice) and compared to control mice (standard housing (SH)-reared mice) (Figure 5A). EE-reared mice exhibited increased locomotion in an open field and earlier eye opening compared to SH-reared pups, indicating developmental advancement (Figure S4).^60,61^ DevATLAS revealed that EE significantly increased the proportion of GCs undergoing circuit maturation in the DG at P20 and P24 (Figures 5B and 5C). Given our earlier findings that tdT-labeled GCs were preferentially utilized to encode contextual memory (Figure 3K and 3L), we hypothesized that the EE-induced precocious tdT labeling in the DG corresponded to improved hippocampus-dependent learning and memory ability.

**Figure 5.**
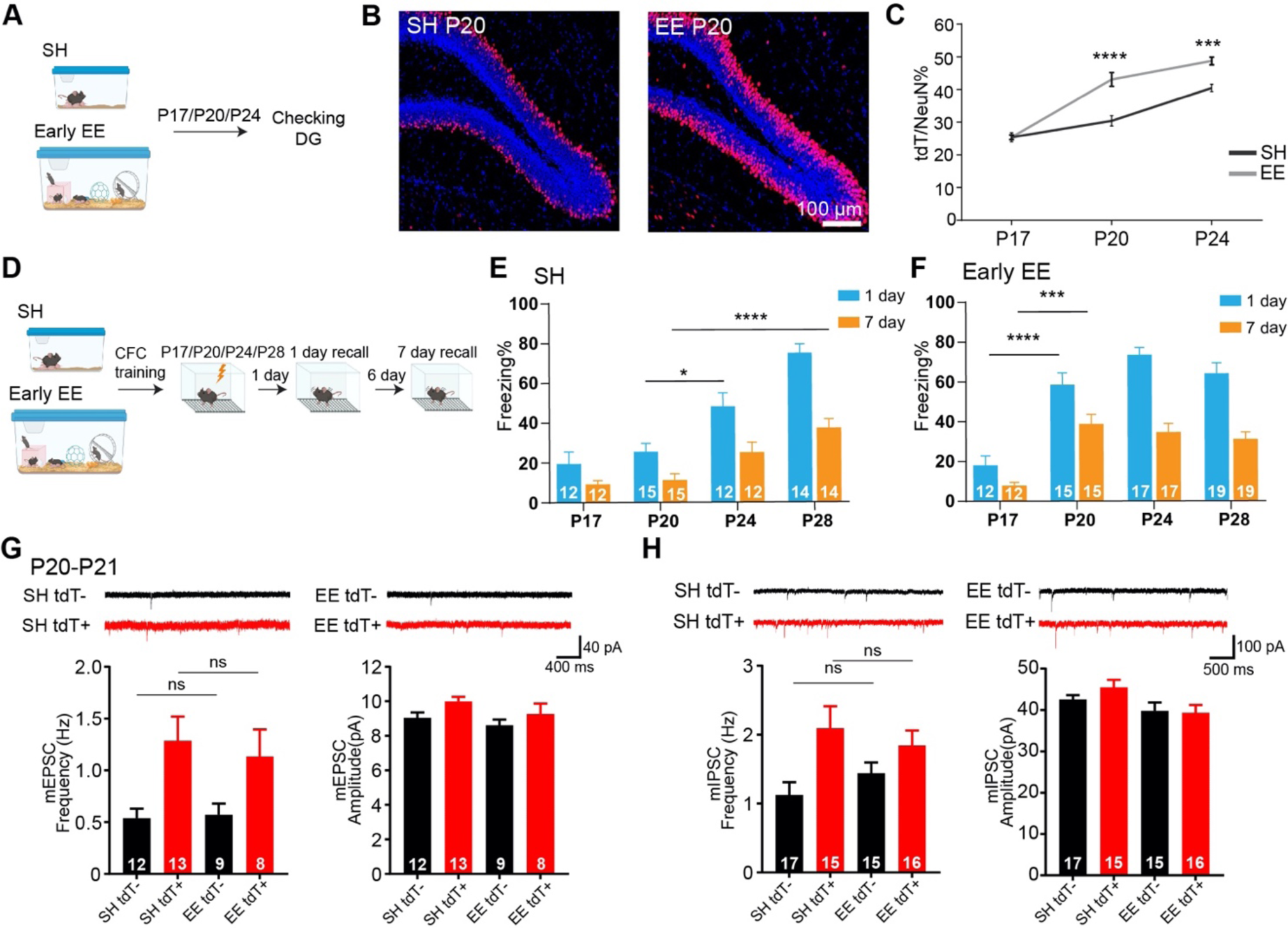
Enriched environment facilitates the circuit maturation in the hippocampus. (A-C) Schematic (A), representative images (B), and quantification (C) of DG tdT signal after standard housed (SH)- and early enriched environment (EE)-rearing. (D) Experimental scheme to test hippocampus-dependent learning capability in SH- and early EE-reared mice. (E) SH-reared mice start displaying significant memory retention starting from P24. (F) EE-reared mice display significant improvement in 1-day and 7-day memory recall by P20. (G and H) mEPSC (G) and mIPSC (H) frequency and amplitude are not different between SH- and EE-reared mice at P20-P21, regardless of tdT labeling. Representative traces are shown on the top. Data are shown as mean ± SEM. *p < 0.05, ***p < 0.001, ****p < 0.0001, ns, not significant. See also Figure S4 and statistical data in Table S1.

We used CFC to evaluate if EE impacted the progress of hippocampus-dependent contextual memory ability during development (Figure 5D, see Methods). Consistent with the literature in rat and mice studies,^13,62^ SH-reared mice younger than P24 displayed little memory recall both one day and seven days after CFC (Figure 5E). Remarkably, EE-reared mice as young as P20 exhibited an ability to retrieve contextual memories after one day and seven days post-CFC, on par with SH-reared P28 mice (Figure 5F).

Critically, the extent of tdT labeling in the DG was highly correlated with the memory ability. At P17, when both EE- and SH-reared mice demonstrated similar memory ability (Figures 5E and 5F), tdT labeling in the DG was unchanged (Figure 5C). However, at P20 and P24, the enhanced memory ability demonstrated by the EE-reared mice corresponded to a significant increase in the proportion of tdT-labeled GCs (Figures 5F and 5C).

Both readouts of DG development (tdT labeling and memory ability) indicate that sensory manipulation influences DG circuit maturation within a specific postnatal window. The sensory experience-induced cognitive advancement was temporary, as by P28, both EE- and SH-reared mice exhibited similar hippocampal-dependent memory ability. Our results demonstrate that sensory engagement can accelerate circuit maturation and cognitive function within a region-specific developmental window.

Interestingly, neither mEPSCs nor mIPSCs of tdT-labeled or unlabeled GCs were significantly changed by EE (Figures 5G and 5H). These data imply that the effect of EE on cognition is due to the abundance of tdT-labeled DG GCs, and that the subsequent synaptic development induced by activation is similar between SH and EE GCs. This is consistent with our finding that tdT-labeled GCs during early development are the ones preferentially recruited to encode contextual memory (Figures 3K and 3L). Taken together, our data consistently indicates that the molecular pathways activated in the tdT-labeled neurons play important roles in activity- and experience-dependent circuit maturation.

### Progressive development of granule cells is linked to activity-dependent transcription programs

To identify the molecular mechanisms underlying *in vivo* circuit maturation, we performed scRNA-seq of the DevATLAS DG at P14, P20, and P24 (Figure 6A), covering the emergence of hippocampus-dependent memory ability (Figure 5E). In total, we obtained 19,047 high-quality single-cell transcriptomes (Figure S5A, see Methods), identifying major populations of neurons, neuronal-fated cells, and glia (Figures 6B, S5B, and S5C). We found that tdT expression was restricted to a subset of mature GCs, as indicated by limited expression of immature markers and increased expression of mature markers (Figures 6C, S5D, and S5E). The proportion of tdT-labeled cells increased with age and matched quantification of tdT labeling in the DG (Figures 5C, 6D, S1F, and S5F).

**Figure 6.**
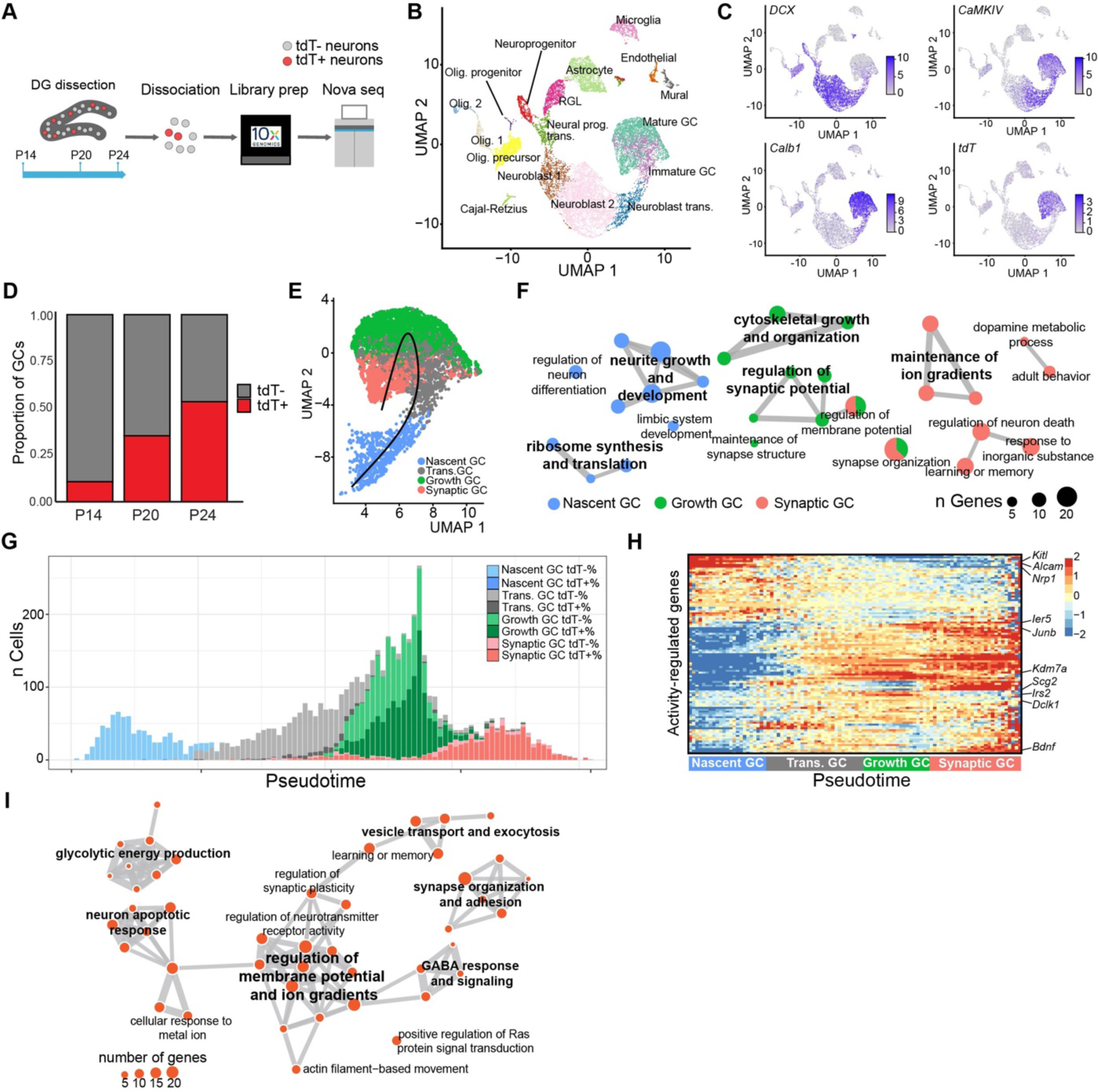
Dentate gyrus granule cell maturation linked to activity-dependent transcription programs. (A) Schematic of DG single-cell sequencing experiment. (B) UMAP plot of single-cell data (n= 19,047 cells) with cell type annotations. GC, granule cell; Trans., transition; prog., progenitor; RGL, radial glia-like cells; Olig., oligodendrocyte. (C) UMAP view of RNA expression of neuronal immature markers (DCX; CaMKIV, Calcium/Calmodulin Dependent Protein Kinase IV), maturation marker (Calb1), and tdT. (D) The proportion of tdT+ GCs at each age reflects previous *in vivo* quantification in Figure 5D. Total cell number: P14, n=1134; P20, n= 2048; P24, n= 2355. tdT+ cell number: P14, n=121; P20, n=722; P24, n=1257. (E) Four clusters of GCs identified using Louvain clustering fall within a single pseudotime trajectory indicated by a black line. Total cell number: Nascent, n=780; Trans., n=1521; Growth, n=2332; Synaptic, n=904. (F) Gene set enrichment analysis (GSEA) identified unique biological contributions of three GC clusters (excluding trans. GC). The proportion of the colors within the node denotes the contribution of genes from each cluster. Edge width reflects the strength of term similarity (based on shared constituent genes). (G) A stacked histogram of GCs across binned pseudotime (100 equally sized bins) indicates that GC clusters reflect progressive developmental and reveals preferential localization of tdT+ GCs towards mature pseudotime endpoint. tdT+ cell number: Nascent, n=2; Trans., n=130; Growth, n=1230; Synaptic, n=873. (H) Activity-regulated gene (ARG) expression across binned pseudotime reveals distinct groups of ARGs are associated with specific GC developmental stages. Synaptic GC population shows the most robust expression of ARGs. (I) GSEA network of DEGs increased in tdT+ GCs across all three ages reveals a tdT phenotype of presynaptic and postsynaptic functional maturity and activation of specific signaling pathways related to Ras, GABA, apoptosis, and energy metabolism. Node size indicates the number of enriched genes contributing to that term. Edge width reflects the strength of term similarity (based on shared constituent genes).

The GC population consisted of two immature and two mature clusters (Figures 6E). Genes involved in neurite extension and protein synthesis were enriched in the immature clusters, while the mature clusters exhibited gene involved in neuronal preparation for synaptic communication (growth GC) and functional synaptic activity (synaptic GC) (Figures 6F and S5G, Table S5). Pseudotime analysis indicated that all GC groups existed in a continuous developmental trajectory (Figure 6E). The increasing representation of tdT-labeled GCs with pseudotime further validated the developmental relevance of the trajectory (Figure 6G). Statistical examination of pseudotime by age and tdT labeling also supported that the tdT-labeled GCs identified in the scRNA-seq dataset were more developmentally advanced than their unlabeled counterparts (Figure S5H). Furthermore, the GC clusters were ordered along pseudotime according to the typical progression of neuronal development (neurite extension, structural reorganization, synaptic signaling) (Figure 6F).

To investigate the link between neuronal activation and the progression of GC development, we examined the expression of activity-regulated genes (ARGs) in the GC population. We isolated a set of ARGs from the literature (see Table S5 for complete list of ARGs used) and characterized their expression across GC pseudotime. We found that many ARGs gradually increased in expression as development progressed, highlighting the importance of activity response in mature GCs (Figure 6H). However, we also identified that separate groups of ARGs were present in different GC clusters (Figures 6H). The ARGs increased in nascent GCs (e.g., *Kitl*, *Alcam*, *Nrp1*) supported the importance of neuronal activity in regulating early neurite outgrowth, migration, pathfinding, and cell adhesion.^63–65^ Some ARGs (e.g., *Kdm7a, Dclk1, Irs2*) were particularly enriched in growth GCs. Importantly, most ARGs and traditional IEGs (*Junb*, *Bdnf*, *Ier5*, and *Scg2*) were robustly expressed within the synaptic GCs (Figure 6H), suggesting that this population is more responsive to activity-dependent stimulus.

Taken together, the characterization of scRNA-seq data from DevATLAS DG reveals possible transcriptional substrates directing activity-dependent circuit development in the DG towards active synaptic signaling. DevATLAS provides us the opportunity to investigate molecular mechanisms which distinguish tdT-labeled GCs from their unlabeled counterparts.

### Energy production pathway is critical for neuronal maturation

In order to identify gene transcripts that may contribute to the development of tdT-labeled GCs, we isolated differentially expressed genes (DEGs) between tdT-labeled and unlabeled GCs at any age (P14, P24, or P24). Gene set enrichment analysis showed that the majority of these DEGs were associated with neuronal maturation, membrane properties, synaptic functions, and, interestingly, glycolytic energy production (Figure 6I). The involvement of energy production pathway in tdT-labeled GCs strongly suggests that the ability to meet the energy demands required for neuronal signaling is highly associated with DG circuit maturation. By extension, our data suggests these processes are critical for cognitive development.

## DISCUSSION

We establish DevATLAS as a novel tool to monitor the sequence of *in vivo* circuit maturation during the early postnatal period (Figures 1, 2, and 4). Our data indicate that circuit assembly is a pre-requisite for DevATLAS activation (Figure 2), that the neuronal activation captured by DevATLAS alters gene expression to confer advanced synaptic function (Figures 3), and that neuronal populations tagged by DevATLAS functionally contribute to behavioral maturity, as seen in the hippocampus (Figures 3 and 5). DevATLAS reveals that brain-wide circuit maturation begins in areas required for basic survival, followed by sensory and motor systems, and lastly cognition-related circuits (Figure 1). Moreover, we constructed the first spatiotemporal map of postnatal whole-brain circuit maturation (Figure 4). This provides us with a powerful platform to identify when and where neural circuit development is perturbed in NDD models, as we demonstrated in our examination of mice with prenatal VPA exposure (Figure 4). Critically, we demonstrated that DevATLAS is also powerful in uncovering molecular mechanisms that facilitate activity-dependent circuit maturation (Figures 6).

### An effective strategy to capture activity-dependent circuit maturation

Compared to other IEG-based genetic and viral tools that label activated neurons,^32,66–69^ DevATLAS specifically captures neurons undergoing activity-dependent circuit maturation during early development. DevATLAS’s unique use of Npas4 induction to tag neurons ensures that the reporter is selectively activity-dependent and neuronal specific. This feature is not present in many other IEG-based systems, as we showed with Fos (Figure S1C). DevATLAS is also unique in capturing neurons that are synaptically and functionally more mature within heterogenous neuronal populations (Figure 3). Npas4’s established role in activity-dependent synaptic development and maintenance of E/I balance may be what positions DevATLAS at the transition from circuit assembly to circuit maturity.^28,30,31,70^ Since different IEGs are involved in specific biological functions,^68^ reporters based on other IEGs, such as Arc,^71^ might provide additional insights into activity-dependent brain development.

DevATLAS tdT labeling accumulates over time. This feature is ideal for whole-brain mapping during development, as it allows to track the timing and abundance of circuit activation in different brain regions. For our purpose, we intentionally avoided CreER for neonatal tdT labeling due to tamoxifen toxicity, which is not well tolerated by embryos and neonates at the concentrations used for maximal labeling, especially in long-term schemas required to achieve longitudinal labeling.^72,73^ Nevertheless, we also made a publicly available Npas4-CreER mouse line (see methods) that allows for monitoring Npas4 induction within experimentally designed time windows.

### A powerful discovery platform for studying NDDs

NDDs typically manifest as a variety of behavioral symptoms by perturbing circuit function without significant alteration of brain structures.^74–76^ Systematically determining when and where circuit function is perturbed is challenging yet necessary to develop effective interventions for patients with NDDs. DevATLAS and the whole-brain imaging analysis pipeline were designed with these goals in mind, offering a fresh perspective by focusing on activity-dependent circuit maturation on the whole-brain scale. Evidences suggest that sensory dysfunction in ASD arises early in development and is attributable, in part, to basal sensory processing regions.^77^ Our findings indicate that layer VIa neurons may play a pivotal role in establishing sensory perception circuits that are disturbed in the prenatal VPA exposure, and possibly in other NDD models as well (Figures 4G-4J). Our data also resonates with a recent finding that reduced spontaneous activity and abnormal development of deep-layer excitatory projection neurons contribute to human NDDs.^78^ Further investigation is needed to reproduce and dissect this phenotype.

### The driving force of activity-dependent circuit maturation

Spontaneous and sensory-driven activity are known to be critical for elements of sensory circuit development like the expansion and refinement of axonal and dendritic arbors.^79,80^ Our study provides compelling evidence that both spontaneous and sensory-driven neuronal activity play prominent roles in early circuit maturation (Figures 1J-1O and S1L-S1P). Although both spontaneous and sensory-driven activity are known to be critical for circuit development, this study provides an effective way to investigate the relative contributions of each type of activity to circuit development, and the timing of the effects varies across the regions. We also demonstrate that early sensory-driven activity can accelerate the maturation and function of non-sensory circuits like the hippocampus (Figure 5). Interestingly, our system does not capture the early stage of global spontaneous activity mediated by the gap-junction,^36,81,82^ suggesting that gap-junction-mediated depolarization may not engage IEG programs that promote synapse formation and remodeling. Our data also highlight the crucial role for glutamatergic transmission in relaying spontaneous activity and sensory input that are responsible for early circuit activation (Figures 1K, 1M, and 2G-2I).^83^

Although the influence of neuromodulators on synaptic development is well-documented,^84,85^ very little is known about how acetylcholine, dopamine, and serotonin influence Npas4 expression. DevATLAS will facilitate future investigations across brain regions to determine the extent and mode by which neuromodulators impact circuit activity during development.

### Cellular and molecular mechanisms of activity-dependent circuit development

Due to the stochastic nature of *in vivo* physiological neuronal activity, previous studies have primarily focused on acute transcription responses triggered by strong, often artificially induced synchronous activity to produce populations of activated neurons *in vitro* or *in vivo*.^86,87^ In contrast, DevATLAS cumulatively captures previously activated neurons, enabling us to uncover molecular and cellular mechanisms that are critical for activity- and experience-dependent circuits maturation under *in vivo* physiological conditions (Figures 5 and 6). The approach we developed here can also be applied to investigate activity-dependent circuit maturation mechanisms across brain regions, cell types, and developmental stages.

## Supporting information

Statistical Table

Whole-brain imaging quantification

Filtered brain regions for hierarchical clustering

VPA whole-brain imaging comparison

Gene expression information for single-cell analysis

## ACKNOWLEDGEMENTS

We thank Xiaochen Sun and Scott J. Neal for critical reading of the manuscript; Xubo Niu, Liu Yang, and Minggui Chen, for the comments of the manuscript; Meizhen Meng, Thi Minh Hieu Lam, and Joslyn Doupe for the suggestions of this study; Martin Hemberg for help with scRNA-seq data analysis; Karen Gentile at the SUNY molecular analysis core for the supports of sequencing experiments; John M. Ashton and Rochester Genomics Center for the scRNA-seq service. This work was funded by Human Frontier Science Program long-term fellowship LT000479/2016-L (J.X.); Brain Research Foundation Seed Grant (Y.L.); Simons Center for the Social Brain Equipment Grant (Y.L.); Paul and Lilah Newton Brain Science Award (Y.L.); NIH grant RF1MH124605 (Y.K.); NIH grant DC014701 (R.Y. & Y.L.); NIH grants NS123710, NS115543, and MH116673 (Y.L.).

## AUTHOR CONTRIBUTIONS

Y.L. conceived and supervised the study. Y.C. and Y.L. made the Npas4-Cre mouse. J.X. and Y.L. designed the experiments. J.X. performed or participated in most of the experiments. A.B. participated in most of the experiments and performed imaging quantification, VPA model, and EE-related experiments. J.T. conducted scRNA-seq data analysis and participated in statistical data analysis. T.Y. conducted the electrophysiology experiments. K.N. and Y.K. established the young mouse brain templates and whole-brain imaging analysis pipeline, and analyzed the whole-brain imaging data. Y.A. established the scRNA-seq analysis pipeline. Q.Q. and R.Y. conducted the naris occlusion experiments. J.X, J.T., and Y.L. wrote the manuscript with inputs from all the authors.

## DECLARATION OF INTERESTS

The authors declare no competing interests.

## SUPPLEMENTARY FIGURES

**Figure S1.**
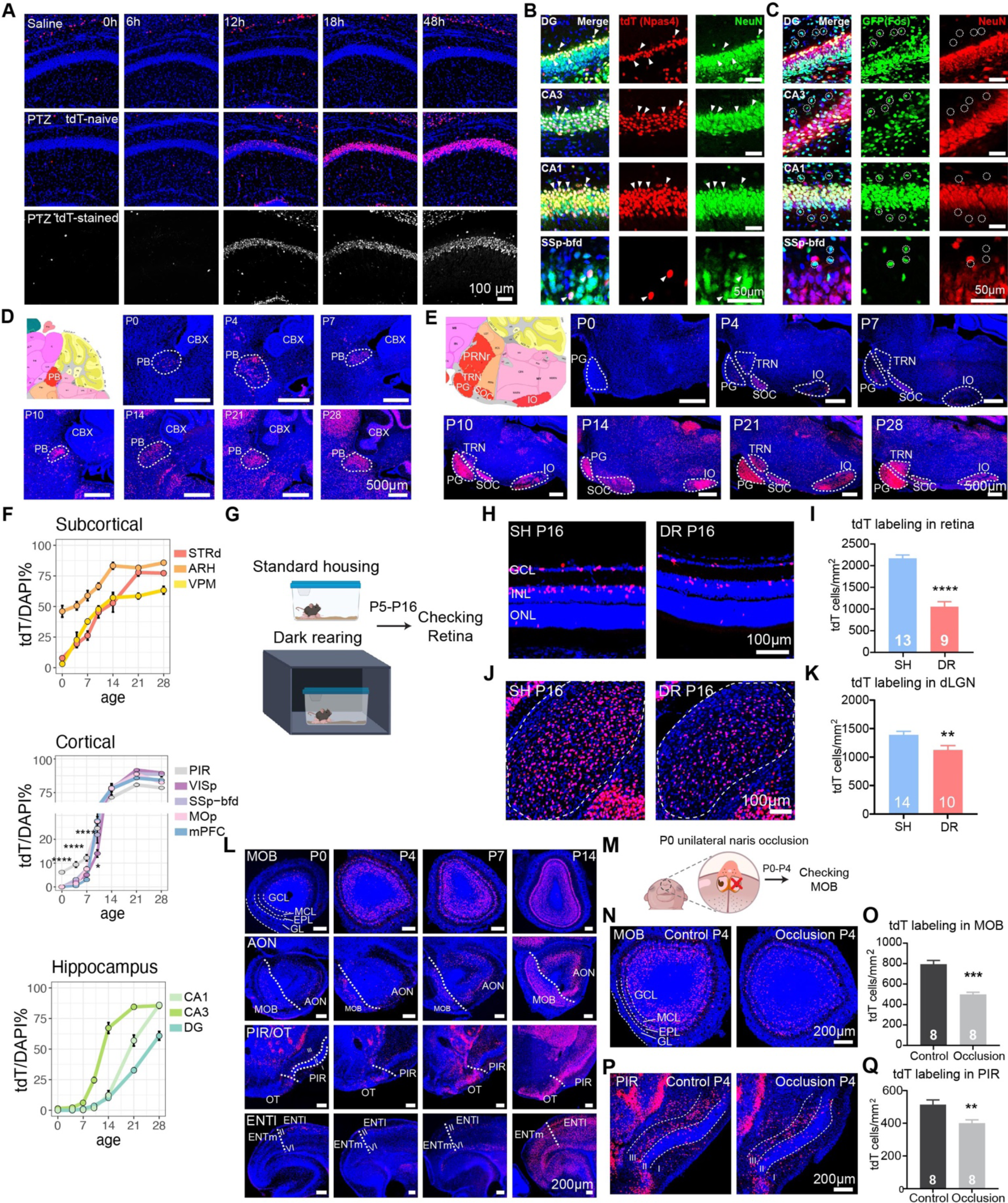
Characterization of tdT labeling in the DevATLAS. (A) Representative images showing tdT labeling in CA1 0h, 6h, 12h, 18h, and 48h after injections of saline and PTZ at P7 in DevATLAS (Npas4-Cre;Ai75) mice. The bottom panel shows tdT antibody staining in the PTZ condition. (B) tdT labeling is neuron specific. Representative images of brain slices from seizure-induced DevATLAS (Npas4-Cre;Ai75) mice (P7) stained with a neuronal marker (NeuN) at P9. The DG, CA3, CA1, and superficial layers of SSp-bfd are shown in the panels, with NeuN staining shown in green. White arrows indicate the overlapping signals between NeuN staining and tdT. (C) Fos activation is not neuronal specific. Representative images of NeuN staining (in red) using P7 brain slices from home cage Fos-tTA;TRE-H2BGFP mice. Dash circles indicate the non-overlapped signals between NeuN staining and tdT signal. (D and E) tdT labeling in the hindbrain from P0 to P28 with anatomical reference to Allen adult mouse brain atlas. PB, parabrachial nucleus; SOC, superior olivary complex; PG, pontine gray; IO, inferior olivary complex; TRN, tegmental reticular nucleus; CBX, cerebellar cortex. (F) Quantification of tdT labeling across 11 brain regions grouped into subcortical, cortical, and hippocampus regions at seven developmental ages. Data is replotted from Figure 1I. (G) Experimental scheme of standard housing (SH) and dark rearing (DR). (H and I) Representative images (H) and quantification (I) showing decreased tdT labeling in retinas in dark-reared mice at P16. (J and K) Representative images (J) and quantification (K) demonstrating decreased tdT labeling in the dLGN of the DR-reared mice compared to SH-reared mice at P16. (L) tdT labeling in olfactory pathway associated regions from P0 to P14 mice. MOB, main olfactory bulb; AON, anterior olfactory nucleus; PIR, piriform cortex; OT, olfactory tubercle; ENTl, entorhinal area, lateral part; ENTm, entorhinal area, medial part; GL, glomerular layer; EPL, external plexiform layer; MCL, mitral cell layer; GCL, granule cell layer. (M) Schematic showing the unilateral naris occlusion experiments. (N and O) Representative images (N) and quantification (O) showing the reduced tdT labeling in MOB after naris occlusion. (P and Q) Representative images (P) and quantification (Q) showing the reduced tdT labeling in PIR after naris occlusion. Data are shown as mean ± SEM. *p < 0.05, **p < 0.01, ***p < 0.001, ****p < 0.0001. See statistical data in Table S1.

**Figure S2.**
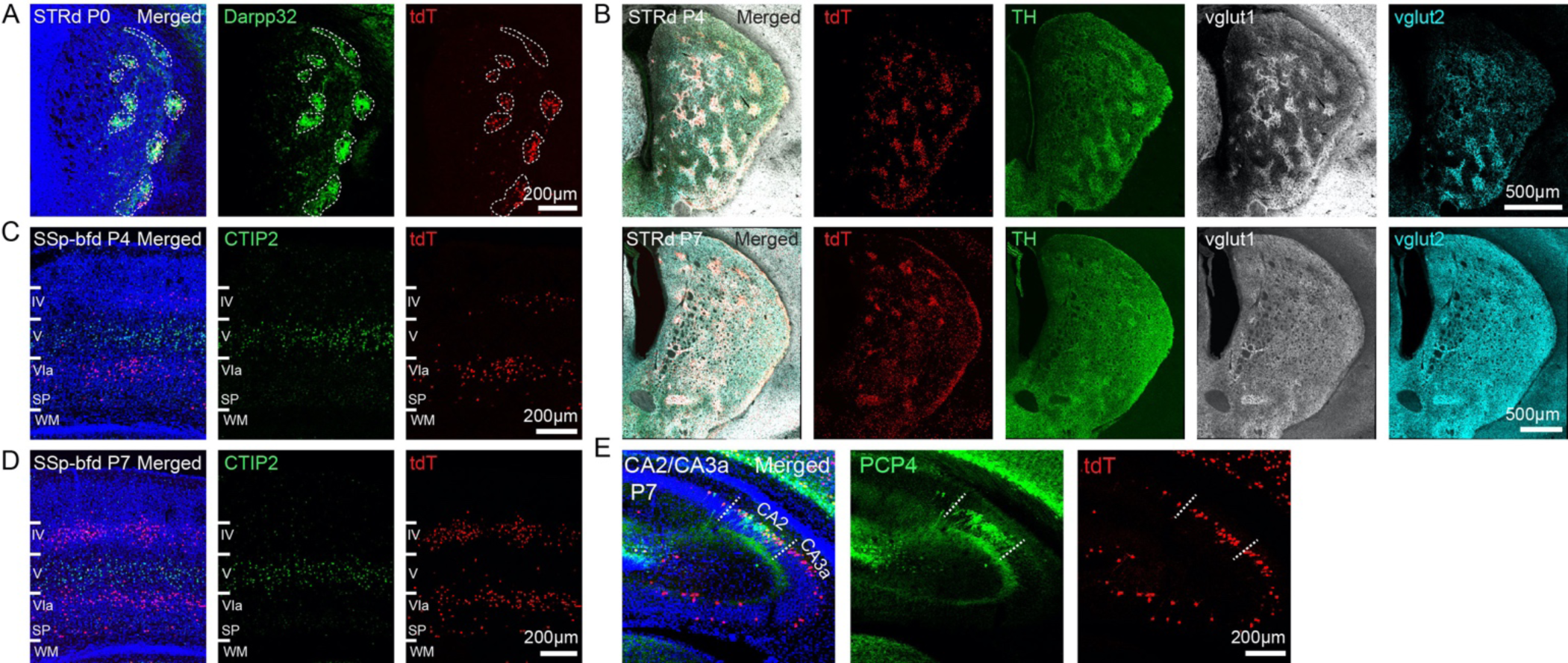
Identify the circuit maturation using markers staining. (A) tdT-labeled regions co-localized with striosome in STRd at P0, indicated by a neonatal striosome marker (Darpp32). (B) Presynaptic markers showed labeling in matrix region of STRd at P7. (C and D) Layer V marker (CTIP2) shows SSp-bfd tdT labeling initiated at layer VI, followed by layer IV at P4 (C) and P7 (D). (E) CA2 (indicated by CA2 marker PCP4) and CA3a exhibited tdT labeling at P7.

**Figure S3.**
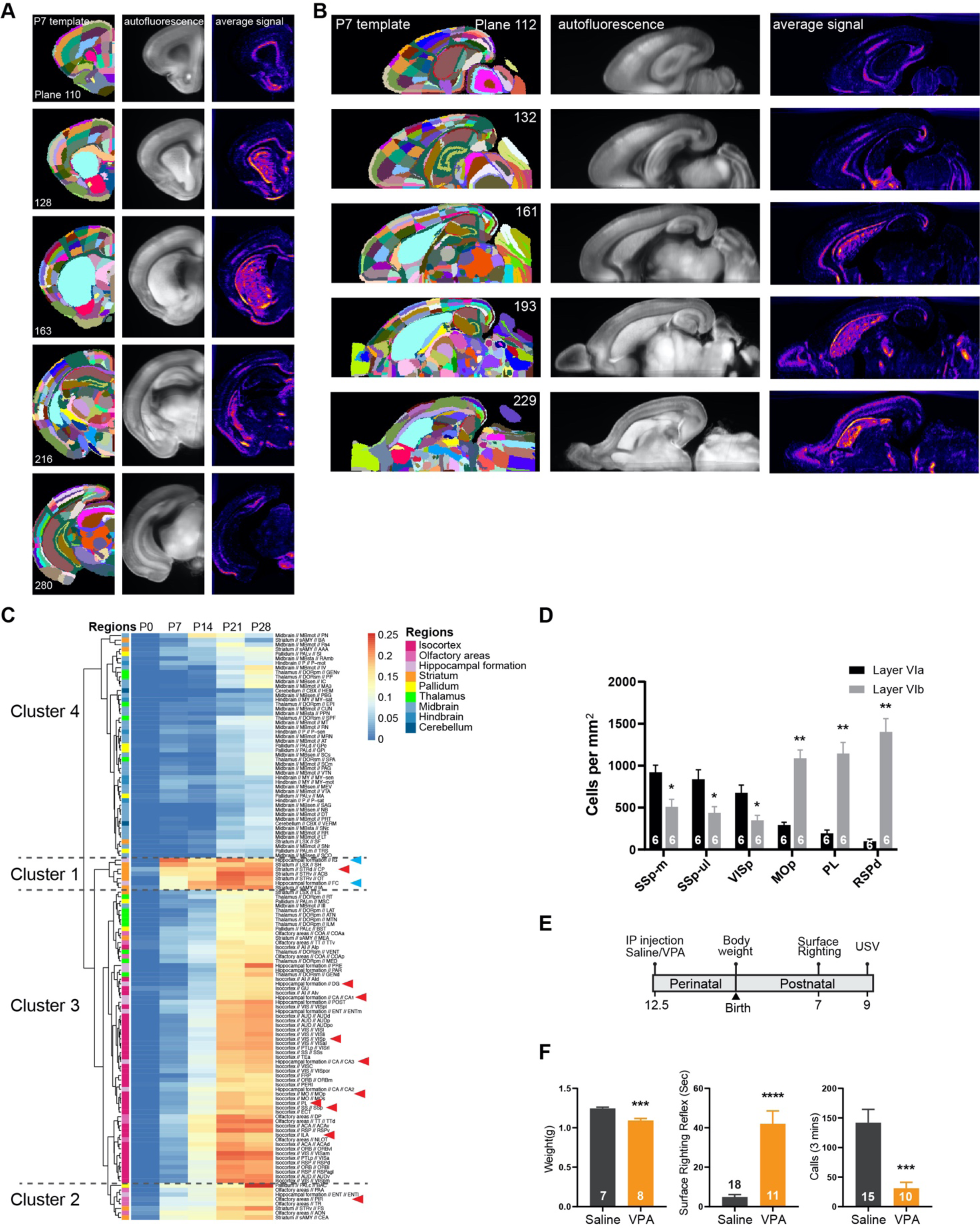
Whole-brain imaging dataset navigation, clustering, and VPA model validation. (A and B) Dataset navigation in coronal (A) and horizontal (B) views with the synchronized template registration, autofluorescence and average tdT signal. (C) Four clusters are identified in hierarchical clustering based on the averaged tdT density in each region from the brains of P0, P7, P14, P21, and P28 mice. Brain regions are color-coded and shown between the dendrogram and heatmap. The regions matching those manually quantified in Figure 1I are marked with red arrows. The fasciola cincerea and induseum griseum are marked with blue arrows. (D) Quantification of the distinct tdT labeling patterns between layer VIa and VIb across different cortical regions at P7. (E) Timeline of saline and VPA injection, body weight measurement and behavioral testing. (F) VPA-treated mice, compared to the saline group, showed decreased body weight at P0, increased surface righting reflex at P7, and a decrease in ultrasonic vocalization at P9. Data are shown as mean ± SEM. *p < 0.05, **p < 0.01, ***p < 0.001, ****p < 0.0001. See statistical data in Table S1.

**Figure S4.**
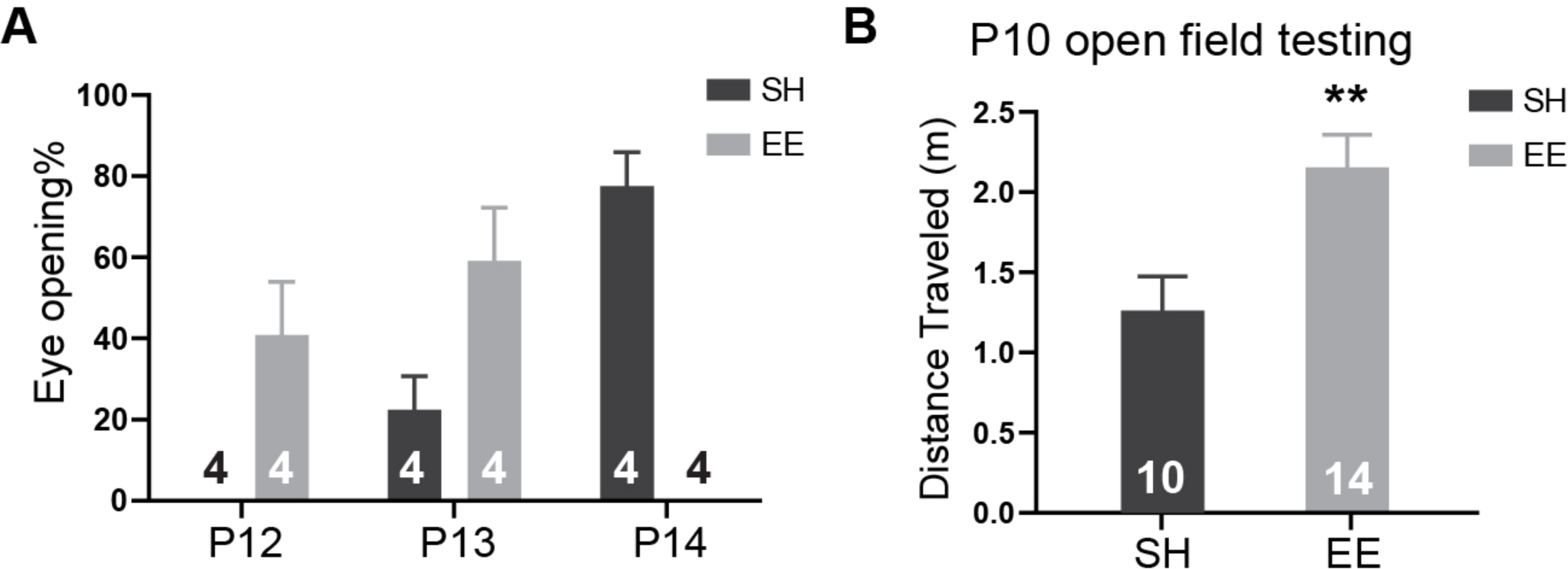
Behavior validation between SH- and EE-reared mice. (A) EE-reared mice open their eyes earlier than SH-reared mice. (B) EE-reared mice exhibit greater locomotion compared to SH-reared mice at P10. Data are shown as mean ± SEM. **p < 0.01. See statistical data in Table S1.

**Figure S5.**
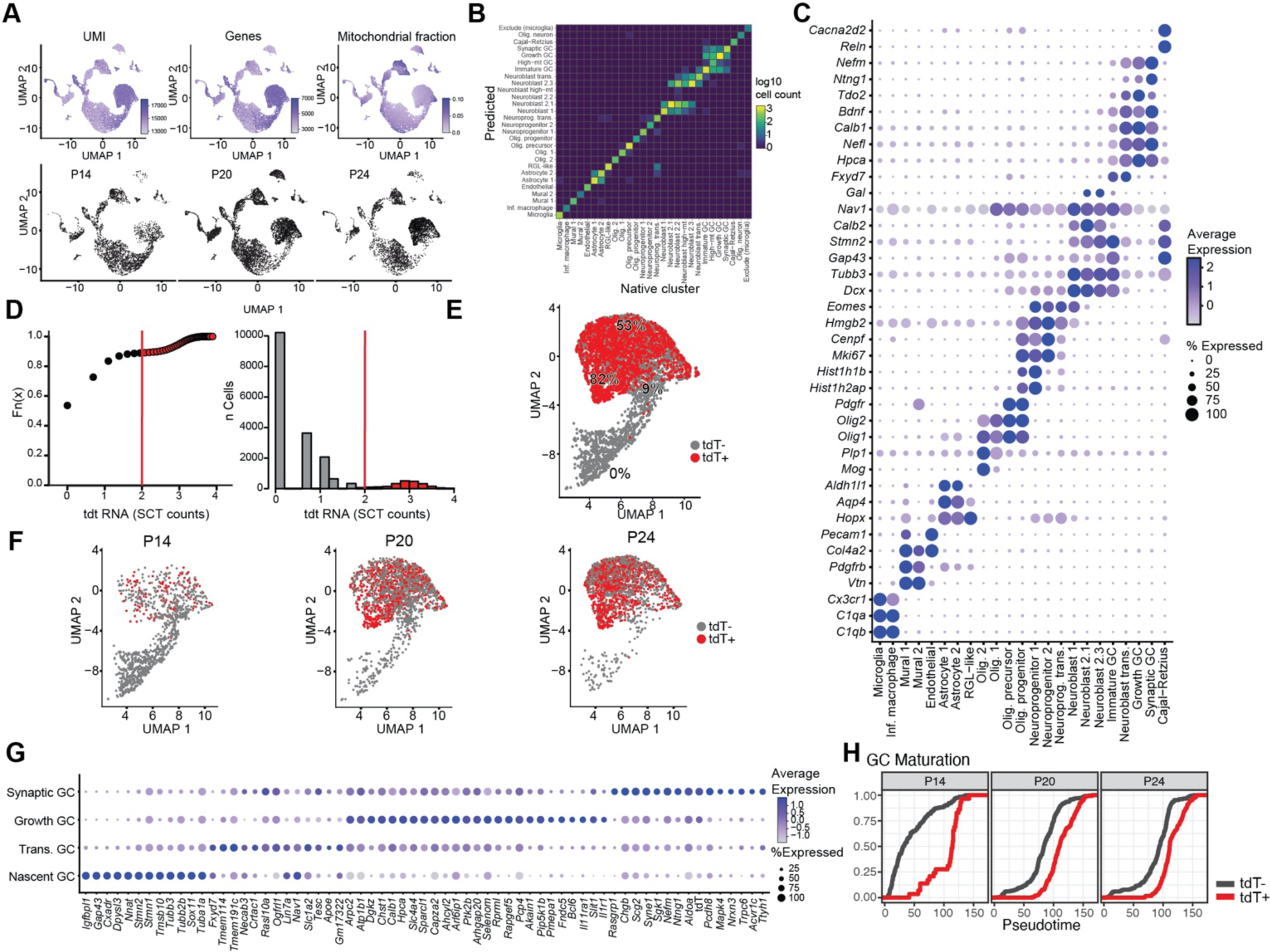
Identification of clusters and tdT status in single-cell data. (A) UMAP plotting showing the number of UMI (unique molecule identifier), genes, mitochondrial fraction, and age (P14, P20, P24) information for each cell. Mean ± SD UMI (raw RNA) = 18,049±9,584, mean ± SD number of genes (raw RNA) = 4,721±1,205, mean ± SD mitochondrial fraction = 0.037±0.016. (B) Cell number plotting of projected cell clusters and native cell clusters to validate cluster consistency. (C) Expression of cell type markers across annotated clusters. (D) tdT status threshold for each cell set as 2 counts after SCTransform. Cell number fractions (Fn) were plotted based on the tdT counts (left), and cell numbers were plotted based on the tdT counts (right). The red line shows the threshold setting. (E) The proportion of tdT-labeled GCs in each subclusters. Nascent tdT+ n=2, Trans tdT+=130, Growth tdT+ n=1230, Synaptic tdT+ n=738. (F) UMAP plotting showing tdT status of cell samples from different ages. (G) Expression of top GC cluster markers. (H) Across all ages, tdT+ GCs display a more mature developmental status than tdT-GCs according to pseudotime analysis. The proportion of population is shown on the y-axis. P14, p<1e-8; P20, p<2.2e-16; P24, p<2.2e-16.

## TABLES

Table S1. Statistical Table

Table S2. Whole-brain imaging quantification

Table S3. Filtered brain regions for hierarchical clustering

Table S4. VPA whole-brain imaging comparison

Table S5. Gene expression information for single-cell analysis

## KEY RESOURCES TABLE

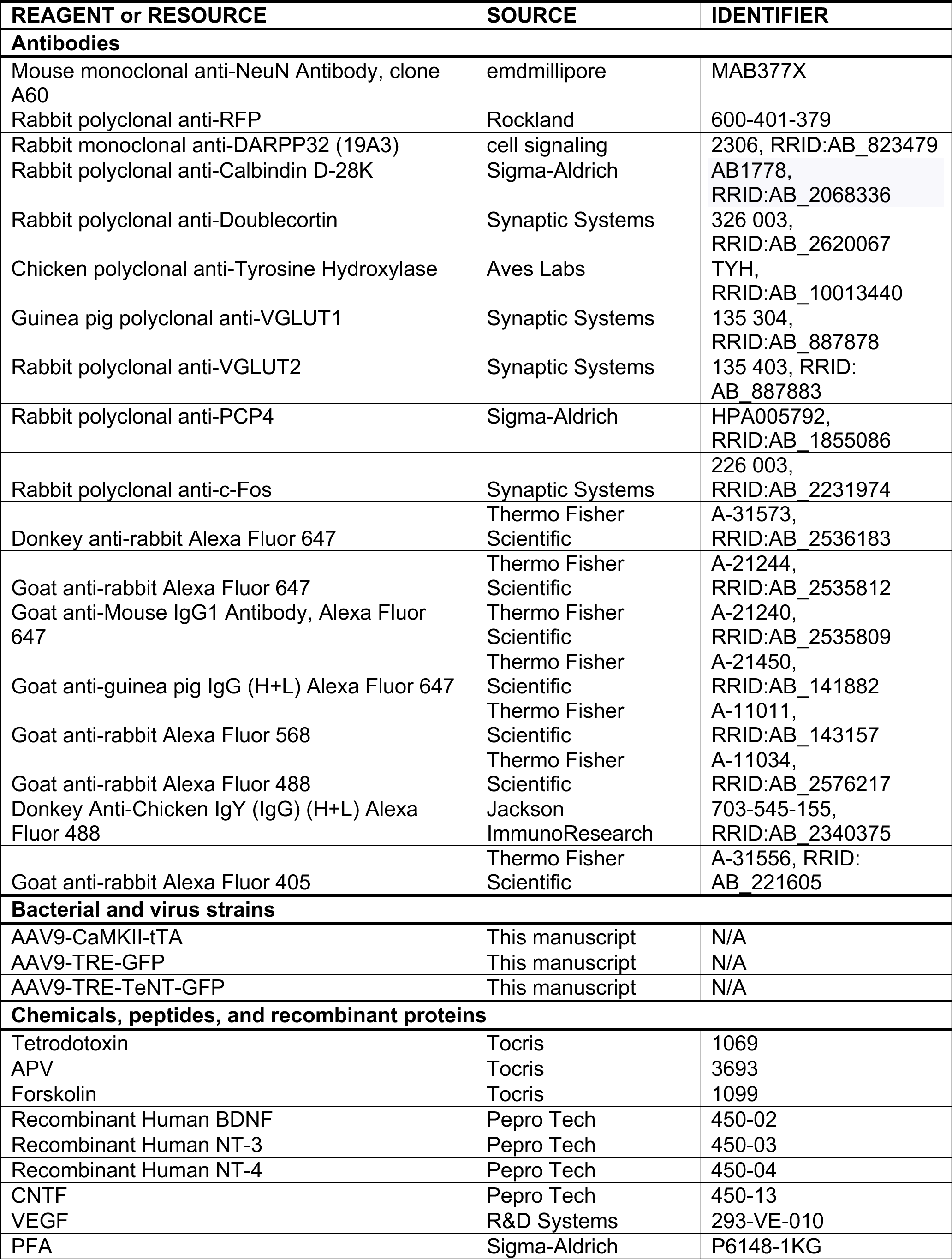
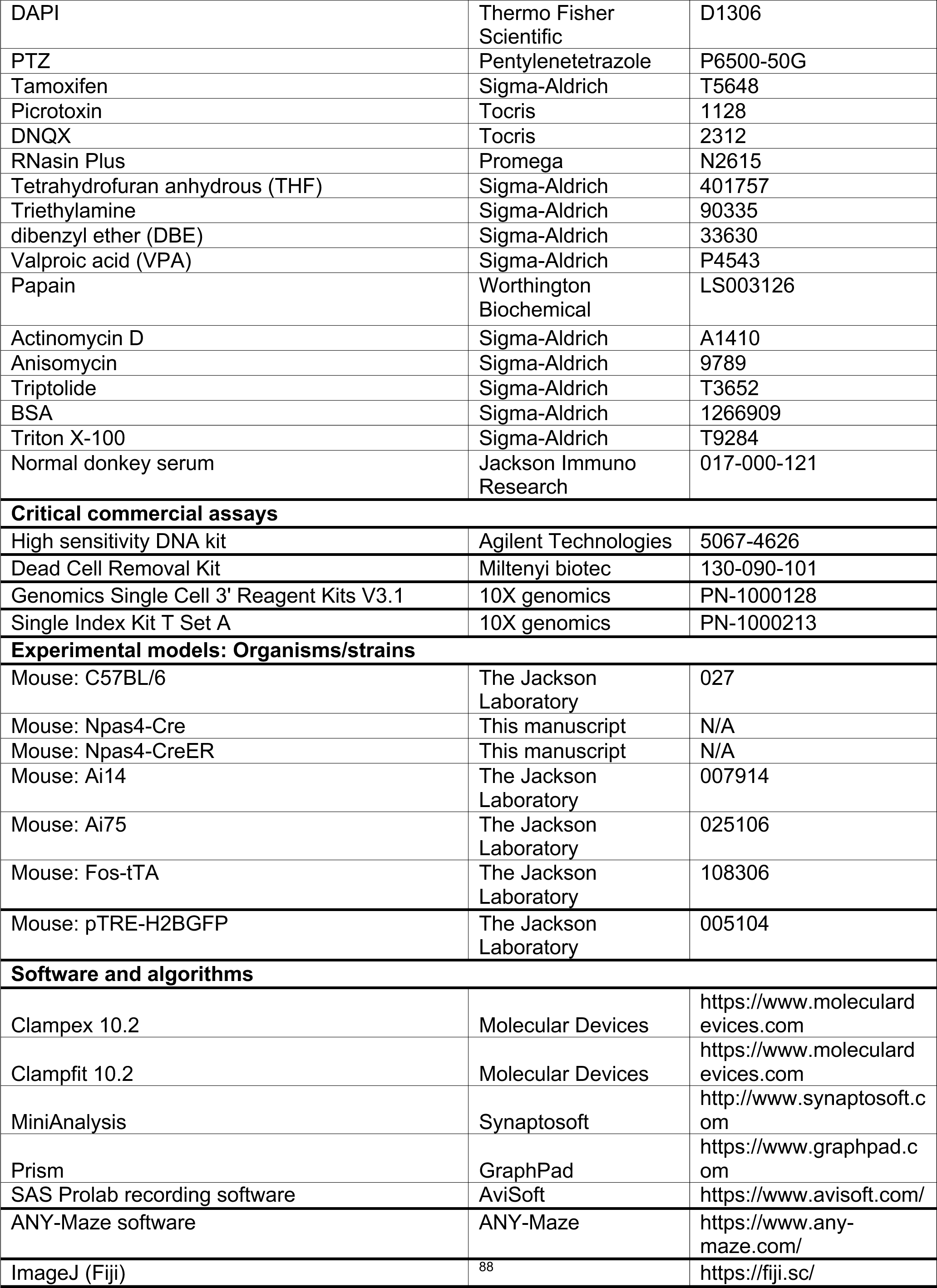
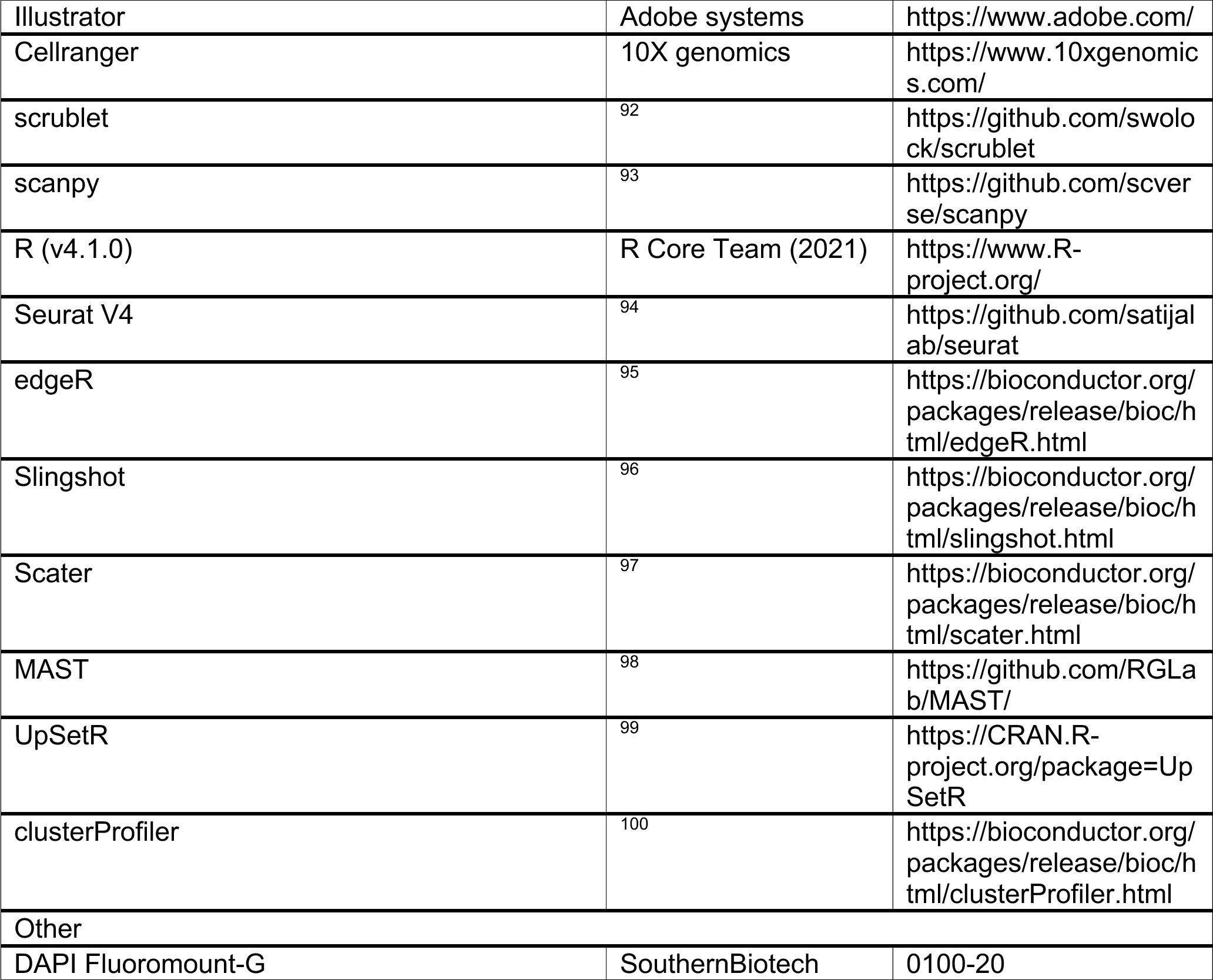

| **REAGENT or RESOURCE** | **SOURCE** | **IDENTIFIER** |
| --- | --- | --- |
| **Antibodies** |  |  |
| Mouse monoclonal anti-NeuN Antibody, clone A60 | emdmillipore | MAB377X |
| Rabbit polyclonal anti-RFP | Rockland | 600-401-379 |
| Rabbit monoclonal anti-DARPP32 (19A3) | cell signaling | 2306, RRID:AB_823479 |
| Rabbit polyclonal anti-Calbindin D-28K | Sigma-Aldrich | AB1778, RRID:AB_2068336 |
| Rabbit polyclonal anti-Doublecortin | Synaptic Systems | 326 003, RRID:AB_2620067 |
| Chicken polyclonal anti-Tyrosine Hydroxylase | Aves Labs | TYH, RRID:AB_10013440 |
| Guinea pig polyclonal anti-VGLUT1 | Synaptic Systems | 135 304, RRID:AB_887878 |
| Rabbit polyclonal anti-VGLUT2 | Synaptic Systems | 135 403, RRID:AB_887883 |
| Rabbit polyclonal anti-PCP4 | Sigma-Aldrich | HPA005792, RRID:AB_1855086 |
| Rabbit polyclonal anti-c-Fos | Synaptic Systems | 226 003, RRID:AB_2231974 |
| Donkey anti-rabbit Alexa Fluor 647 | Thermo Fisher Scientific | A-31573, RRID:AB_2536183 |
| Goat anti-rabbit Alexa Fluor 647 | Thermo Fisher Scientific | A-21244, RRID:AB_2535812 |
| Goat anti-Mouse IgG1 Antibody, Alexa Fluor 647 | Thermo Fisher Scientific | A-21240, RRID:AB_2535809 |
| Goat anti-guinea pig IgG (H+L) Alexa Fluor 647 | Thermo Fisher Scientific | A-21450, RRID:AB_141882 |
| Goat anti-rabbit Alexa Fluor 568 | Thermo Fisher Scientific | A-11011, RRID:AB_143157 |
| Goat anti-rabbit Alexa Fluor 488 | Thermo Fisher Scientific | A-11034, RRID:AB_2576217 |
| Donkey Anti-Chicken IgY (IgG) (H+L) Alexa Fluor 488 | Jackson ImmunoResearch | 703-545-155, RRID:AB_2340375 |
| Goat anti-rabbit Alexa Fluor 405 | Thermo Fisher Scientific | A-31556, RRID:AB_221605 |
| **Bacterial and virus strains** |  |  |
| AAV9-CaMKII-tTA | This manuscript | N/A |
| AAV9-TRE-GFP | This manuscript | N/A |
| AAV9-TRE-TeNT-GFP | This manuscript | N/A |
| **Chemicals, peptides, and recombinant proteins** |  |  |
| Tetrodotoxin | Tocris | 1069 |
| APV | Tocris | 3693 |
| Forskolin | Tocris | 1099 |
| Recombinant Human BDNF | Pepro Tech | 450-02 |
| Recombinant Human NT-3 | Pepro Tech | 450-03 |
| Recombinant Human NT-4 | Pepro Tech | 450-04 |
| CNTF | Pepro Tech | 450-13 |
| VEGF | R&D Systems | 293-VE-010 |
| PFA | Sigma-Aldrich | P6148-1KG |
| DAPI | Thermo Fisher Scientific | D1306 |
| PTZ | Pentylentetrazole | P6500-50G |
| Tamoxifen | Sigma-Aldrich | T5648 |
| Picrotoxin | Tocris | 1128 |
| DNQX | Tocris | 2312 |
| RNasin Plus | Promega | N2615 |
| Tetrahydrofuran anhydrous (THF) | Sigma-Aldrich | 401757 |
| Triethylamine | Sigma-Aldrich | 90335 |
| dibenzyl ether (DBE) | Sigma-Aldrich | 33630 |
| Valproic acid (VPA) | Sigma-Aldrich | P4543 |
| Papain | Worthington Biochemical | LS003126 |
| Actinomycin D | Sigma-Aldrich | A1410 |
| Anisomycin | Sigma-Aldrich | 9789 |
| Triptolide | Sigma-Aldrich | T3652 |
| BSA | Sigma-Aldrich | 1266909 |
| Triton X-100 | Sigma-Aldrich | T9284 |
| Normal donkey serum | Jackson Immuno Research | 017-000-121 |
| **Critical commercial assays** |  |  |
| High sensitivity DNA kit | Agilent Technologies | 5067-4626 |
| Dead Cell Removal Kit | Miltenyi biotec | 130-090-101 |
| Genomics Single Cell 3' Reagent Kits V3.1 | 10X genomics | PN-1000128 |
| Single Index Kit T Set A | 10X genomics | PN-1000213 |
| **Experimental models: Organisms/strains** |  |  |
| Mouse: C57BL/6 | The Jackson Laboratory | 027 |
| Mouse: Npas4-Cre | This manuscript | N/A |
| Mouse: Npas4-CreER | This manuscript | N/A |
| Mouse: Ai14 | The Jackson Laboratory | 007914 |
| Mouse: Ai75 | The Jackson Laboratory | 025106 |
| Mouse: Fos-tTA | The Jackson Laboratory | 108306 |
| Mouse: pTRE-H2BGFP | The Jackson Laboratory | 005104 |
| **Software and algorithms** |  |  |
| Clampex 10.2 | Molecular Devices | https://www.moleculardevices.com |
| Clampfit 10.2 | Molecular Devices | https://www.moleculardevices.com |
| MiniAnalysis | Synaptosoft | http://www.synaptosoft.com |
| Prism | GraphPad | https://www.graphpad.com |
| SAS Prolab recording software | AviSoft | https://www.avisoft.com/ |
| ANY-Maze software | ANY-Maze | https://www.any-maze.com/ |
| ImageJ (Fiji) | 88 | https://fiji.sc/ |
| Illustrator | Adobe systems | https://www.adobe.com/ |
| Cellranger | 10X genomics | https://www.10xgenomics.com/ |
| scrublet | <sup>92</sup> | https://github.com/swolock/scrublet |
| scanpy | <sup>93</sup> | https://github.com/scverse/scanpy |
| R (v4.1.0) | R Core Team (2021) | https://www.R-project.org/ |
| Seurat V4 | <sup>94</sup> | https://github.com/satijalab/seurat |
| edgeR | <sup>95</sup> | https://bioconductor.org/packages/release/bioc/html/edgeR.html |
| Slingshot | <sup>96</sup> | https://bioconductor.org/packages/release/bioc/html/slingshot.html |
| Scater | <sup>97</sup> | https://bioconductor.org/packages/release/bioc/html/scater.html |
| MAST | <sup>98</sup> | https://github.com/RGLab/MAST/ |
| UpSetR | <sup>99</sup> | https://CRAN.R-project.org/package=UpSetR |
| clusterProfiler | <sup>100</sup> | https://bioconductor.org/packages/release/bioc/html/clusterProfiler.html |
| Other |  |  |
| DAPI Fluoromount-G | SouthernBiotech | 0100-20 |

### RESOURCE AVAILABILITY

#### Lead contact

Further information and requests for resources and reagents should be directed to the lead contact: Yingxi Lin.

#### Materials availability

Animal lines and datasets generated in this study are available on the request from lead contact.

#### Data and code availability

All original codes will be available on Github.

### EXPERIMENTAL MODEL AND SUBJECT DETAILS

#### Mouse line

C57BL/6 mice were purchased from the Charles River Laboratory. Npas4-Cre and Npas4-CreER were generated in this study. Ai14 (*Gt(ROSA)26Sor^tm^*^14^(CAG–tdTomato)*^Hze^*), Ai75 (Gt(ROSA)26Sor^tm75.1(CAG-tdTomato*)Hze^), Fos-tTA (Tg(Fos-tTA,Fos-EGFP*)1Mmay), pTRE-H2BGFP (Tg(tetO-HIST1H2BJ/GFP)47Efu) were purchased from the Jackson Laboratory. All mice have been backcrossed with wild-type animals from the Charles River Laboratory. All mice were housed in a 12-hour light-dark cycle with unrestricted access to food and water. Mice were weaned on postnatal day 21, unless otherwise specified. Post-weaning mice were housed by sex in groups of 3-5 until used for experiments. As standard practice, both males and females were used in this study. Animal protocols were performed in accordance with NIH guidelines and approved by the IACUCs of SUNY Upstate Medical University or Stowers Institute for Medical Research.

### METHOD DETAILS

#### Cell culture

Cortical primary neuron culture was prepared from Npas4Cre;Ai14 P1 brains according to previous publication.^28^ Neurons were plated on glass slides within 24-well plates with 150k neurons per slide and one slide per well. Neurons were cultured in Neurobasal A medium (NBA, Invitrogen) supplemented with B27 (Invitrogen), GlutaMAX (Invitrogen), TTX (0.5 mM, Tocris), and APV (50 mM, Tocris), and were maintained in a humidified incubator with 5% CO_2_ at 37°C.

On day in vitro (DIV) 4, the medium was removed and reserved, and neurons were stimulated with NBA containing one of the following: KCl (50 mM), KCl+EGTA (5mM), forskolin (10 μM, Tocris), BDNF (50 ng/ml, Pepro Tech), NT3 (50 ng/ml, Pepro Tech), NT4 (50 ng/ml, Pepro Tech), CNTF (100 ng/ml, Pepro Tech), or VEGF (100 ng/ml, R&D Systems). After 1 hour of stimulation, the medium was replaced with reserved conditioned medium. On DIV6, neurons were fixed in 4% paraformaldehyde (PFA) (Sigma-Aldrich) in phosphate buffered saline (PBS) for subsequent imaging and quantification.

#### Viral vectors and titers

All vectors and viruses were prepared in house. To block the synaptic transmission from the entorhinal cortex to the dentate gyrus, AAV9-CaMKII-tTA was co-injected with AAV9-TRE-GFP or AAV9-TRE-TeNT-GFP into the medial entorhinal cortex and the lateral entorhinal cortex of postnatal 1-day (P1) mice. Examination of tdT labeling was carried out at P16.

#### Stereotaxic Injection

Animals were gas-anesthetized using 1.5% isoflurane in O_2_ and received bi-lateral injections in the medial entorhinal cortex and the lateral entorhinal cortex. The volume for injection was 36nl for each side. The surgery animals were returned to their dam and sacrificed according to the procedure.

#### Euthanasia and tissue processing for fluorescent imaging

Npas4-Cre;Ai75 animals were used for all quantifications of tdT-labeled cells. Animals were euthanized via exsanguination and were transcardially perfused with PBS followed by 4% PFA in PBS. Brains were post-fixed in 4% PFA at 4°C overnight and then incubated in 30% sucrose at 4°C for 24 hours. Brains were sectioned at 50 um using a cryostat (Leica). Corresponding brain slices were washed 3 times in PBS.

To quantify tdT-labeled cells without any additional antibody staining, tissue slices were stained with DAPI (1 μg/ml, Sigma-Aldrich) for 10 min, washed three times with PBS, and mounted on microscope slides.

#### Euthanasia and tissue processing for electrophysiology

Mice were anesthetized with isoflurane (P14 and P20) or on the ice (P7) and decapitated. Brains were then immediately dissected out and cooled in ice-cold cutting solution containing 210 mM sucrose, 2.5 mM KCl, 1.24 mM NaH_2_PO_4_, 8 mM MgCl_2_, 1 mM CaCl_2_, 26 mM NaHCO_3_, 20 mM Glucose, and 1.3 mM sodium-L-ascorbate, with ∼340 mOsm osmolarity and pH of 7.3. After a few minutes, the brains were cut into 300 μM transverse slices using a vibratome (VT1200, Leica). Slices were then immediately transferred to a warm recovery solution (32°C) with 50% cutting solution and 50% ACSF. ACSF contains 119 mM NaCl, 2.5 mM KCl, 1.24 mM NaH_2_PO_4_, 1.3 mM MgCl_2_, 2.5 mM CaCl_2_, 26 mM NaHCO_3_, and 10 mM glucose, with 305 mOsm osmolarity and pH of 7.3. After 12 minutes of recovery, the slices were transferred to ACSF (∼25°C) and bubbled with carbogen gas (95% O_2_ and 5% CO_2_) for at least 30 minutes at room temperature (RT).

#### Euthanasia and tissue processing for RNA sequencing

For scRNA-seq, a similar approach was employed when collecting brain samples. Artificial cerebrospinal fluid (ACSF) was prepared prior to collecting the samples and was modified according to a previous publication.^86^ ACSF contains 124 mM NaCl, 2.5 mM KCl, 1.2 mM NaH_2_PO_4_, 24 mM NaHCO_3_, 5 mM HEPES, 13 mM glucose, 2 mM MgSO_4_, and 2 mM CaCl_2_. A cocktail of chemicals was added into ACSF, including 1 μM TTX (Sigma), 100 μM APV (Tocris), 1 μM DNQX (Tocris), 5 μg actinomycin D (Sigma) per mL, 10 μg anisomycin (Sigma) per ml, and 10 μM triptolide (Sigma). ACSF was bubbled with carbogen gas (95% O_2_ and 5% CO_2_) for at least 15 minutes prior to use.

Mice were anesthetized with isoflurane (P14, P20, P24) for 5 minutes and then euthanized by exsanguination and perfused with ice-cold ASCF. The brain was rapidly dissected and sliced into 400 μm sections using a vibratome, with ACSF used as the cutting solution. Brain slices were then processed according to the single-cell protocol.

#### Immunohistochemistry

After three washes in PBS, brain slices were incubated in blocking buffer containing 10% goat serum and 1% Triton-X100 (Sigma-Aldrich) in PBS at RT for 2 hours. Subsequently, the sections were incubated with primary antibodies in staining buffer (5% goat serum and 0.1% Triton-X100 in PBS) at 4°C overnight. After primary incubation, the sections were washed three more times in PBS and then were incubated with secondary antibodies in 5% goat serum and 0.1% Triton-X100 in PBS at RT for 2 hours. After a final series of the three PBS washes, the sections were mounted on microscope slides with DAPI Fluoromount-G (SouthernBiotech) and covered with coverslips. The following primary antibodies were used: rabbit polyclonal anti-c-Fos (1:2000, Synaptic System), mouse monoclonal IgG1 anti-NeuN (1:500, emdmillipore), and rabbit polyclonal anti-RFP (1:500, Rockland). The secondary antibodies used were anti-primary species with Alexa Fluor 568 and 647 (1:500, Thermo Fisher Scientific).

#### Brain slice imaging and quantification

For most brain regions, we acquired images with 10X and 20X objective lens using image stitching. For the dentate gyrus (DG), we acquired images with an 60X objective lens due to high cell density. Manual image quantification was performed using ImageJ software. Images were cropped, annotated, and arranged using Adobe Illustrator software (Adobe systems).

#### Inducing postnatal seizures

To induce seizure, P7 animals were intraperitoneally (IP) injected with PTZ (dissolved in sterile saline, ∼7.2 of pH) with a dosage of 100 mg/kg. As a control, littermates were injected with saline. Animals were euthanized and perfused at 0h, 6h, 12h, 18h, and 48 h (hours) after injection. Brains were post-fixed with 4% PFA at 4°C overnight.

#### Visual-dependent sensory experiments

To examine the role of retinal activity on circuit development, unilateral or bilateral enucleation was performed on P1 pups. All surgical tools were sterilized with 70% ethanol. Animals were injected with buprenorphine ER (0.04mg/kg) 10 minutes before the procedure and then anesthetized on ice for one minute. After verifying the extent of anesthesia with a toe-pinch, the skin around the eyes was sterilized with 70% ethanol. The eyelids were then opened with a scalpel to expose the eye which was then carefully excised from the orbit with forceps and scissors under a dissection scope. Following eye removal, cotton tips were used to soak up blood and provide light pressure on the orbit to stop bleeding. The eyelid was closed with forceps and sealed using approximately 5 μl of Vetbond glue (3M, MN, USA). After the surgery, animals were transferred to a heating pad for recovery. During recovery, a thin coat of lidocaine hydrochloride jelly USP, 2% (Akorn, Lake Forest, IL, USA) was applied to the eyelid to prevent pain. Once recovered, the pups were returned to the home cage with the dam. Brain samples were collected according to the experiment design.

To examine the impact of spontaneous retinal activity on circuit development, a cohort of animals were raised in a light-tight container in a dark room from P5 to P16. Control animals for dark rearing experiment were raised under 12h:12h light-dark cycle in standard animal facility conditions. Brain samples were collected at P10 or P16 according to the experimental design. To validate the light deprivation, an X-ray film was placed in the same box and developed at the end of deprivation to indicate the presence of light contamination.

#### Olfactory deprivation: unilateral naris occlusion

Unilateral naris occlusion was performed at P0. Pups were anesthetized on ice for one minute. After verifying the extent of anesthesia with a toe-pinch, animals were immediately subjected to electrocautery for ∼2s on left or right side of nostril under a dissection scope. During the procedures, any contact of skin surface with electrocautery device was avoided. Pups were then recovered on a heat pad until they woke up and transferred to the dam. The nose were immersed in water daily to check for air bubbles to ensure nasal blockage of the cauterized nostril. Brain samples were collected at P4.

#### Electrophysiology

Electrophysiological measurements of developing neurons were performed at specific developmental timepoints: striatal measurements were collected at P7, and DG measurements were collected at P14 and P20. All recordings were performed in a system that continuously perfused with carbogenated ACSF at RT with a flow rate of 2 mL/min. Slices were gently placed and fixed in the recording chamber, and tdT-labeled cells were identified using a laser. Borosilicate glass pipettes with 3-6 MU tip resistance were used for recording. Data were collected using a Multiclamp 700B (Molecular Devices), filtered at 3 kHz and digitized at 10 kHz using a Digidata 1440A and Clampex 10.2 software (Molecular Devices). mEPSCs were pharmacologically isolated with 0.5 mM tetrodotoxin (TTX, Tocris) and 50 mM picrotoxin (Tocris) in the ACSF and recorded with voltage clamp mode at −70mV. The internal solution for mEPSC contained 130 mM CsMeSO_3_, 10 mM phosphocreatine, 1 mM MgCl_2_, 10 mM HEPES, 0.2 mM EGTA, 4 mM Mg-ATP, and 0.5 mM Na-GTP, at 295 mOsm osmolarity with pH adjusted to 7.25 using CsOH. mIPSC were pharmacologically isolated with ACSF containing 0.5 mM TTX and 50 mM APV (Tocris) and 20 mM DNQX (Tocris), with cells voltage clamped at −70mV. The high-Cl solution contained 103 mM CsCl, 12 mM CsMeSO_3_, 5 mM TEA-Cl, 10 mM HEPES, 4 mM Mg-ATP, 0.5 mM Na-GTP, 0.5 mM EGTA, and 1 mM MgCl_2_, at 295 mOsm osmolarity with pH adjusted to 7.25 using CsOH.. The resting membrane potential, reversal potential of GABA responses (EGABA), GABA driving force, and cellular Cl-concentration were measured according to previous publications.^101,102^

Data were blindly analyzed using MiniAnalysis (Synaptosoft) and Clampfit 10.2 (Molecular Devices).

#### Single-cell sequencing library preparation

The dissociation buffer was modified according to previous publication.^86^ Dissociation buffer contained HBSS (Life Technologies) with 10 mM HEPES (Sigma-Aldrich), 172 mg kynurenic acid (Sigma-Aldrich) per liter, 0.86 g MgCl_2_·6H2O per liter, and 6.3 g d-glucose per liter, pH adjusted to 7.35. Papain was added to the dissociation solution with a final concentration of 20 U/ml. A cocktail of chemicals was added, including 1 μM TTX (Sigma-Aldrich), 100 μM APV (Tocris), 1 μM DNQX (Tocris), 5 μg actinomycin D (Sigma-Aldrich) per milliliter, 10 μg anisomycin (Sigma-Aldrich) per milliliter, and 10 μM triptolide (Sigma-Aldrich). Dissociation buffer bubbled with carbogen gas (95% O_2_ and 5% CO_2_) for at least 15 minutes prior to use.

Brain slices were collected from P14 (one library), P20 (two libraries), and P24 (one library) for SH reared mice, with two males and two females were contained in each library. The DG was dissected from the brain slices under a dissecting scope. The ACSF used for cutting was then replaced with a dissociation buffer after samples had been collected from all animals. The dissected tissue was trimmed into small pieces and then pipetted up-and-down twice with a 1 ml pipette tip and gently shaken at 37°C for 20 minutes. The dissociation buffer was then replaced with fresh, ice-cold dissociation buffer without papain. The tissue pieces were then moved to a 15 ml Falcon tube and gently triturated with a 1 ml tip up-and-down for 10 times. The suspension was passed through a 40 μm filter to remove clumps.

The filtered cell solution was centrifuged at 180 g for 3 minutes and washed four times with a dissociation buffer containing 0.04% BSA. Unhealthy cells were removed according to dead cell removal kit instructions (Miltenyi biotec). After the last wash, cells were resuspended in 100 μl dissociation buffer containing 0.04% BSA without any blockers. The cell suspension then passed through a 15 μm filter to remove any remaining clumps or debris and kept on ice. Manual cell counting was immediately performed using a hemocytometer, and the volume of cell suspension needed for single-cell library preparation was determined by cell density. Single-cell libraries were then made according to the 10X Genomics Single Cell 3’ Reagent Kits V3.1 User Guide (10X genomics). Libraries were indexed by Single Index Kit T Set A (10X genomics). Multiple libraries were pooled and sequenced with Nova seq at Genomics Research Center of University of Rochester.

#### Whole-brain clearing and imaging

We followed the FDISCO protocol to preserve endogenous tdT fluorescence from the developmental brains of Npas4-Cre;Ai75 mice.^53^ P0, P4, P7 animals were anesthetized on ice and P10, P14, P21, and P28 animals were anesthetized with isoflurane and then euthanized via exsanguination and transcardially perfused with PBS and then 4% PFA in PBS. Brains were post-fixed in 4% PFA at 4°C overnight and then washed in PBS three times for 1 hour each. The subsequent dehydration and clearing steps were performed at 4°C with gentle shaking and without any light exposure. Samples were dehydrated in a series of tetrahydrofuran anhydrous (THF, Sigma-Aldrich) solution concentrations for 12 hours at each concentration (50%, 70%, 80%, and 100% volume). Each solution was diluted in ddH_2_O, and pH adjusted to 9.0 with triethylamine (Sigma-Aldrich). A final step of dehydration was performed using 100% THF for 12 hours. Following dehydration, refractive index matching and clearing were achieved by washing brains twice in dibenzyl ether (DBE, Sigma-Aldrich) for 3 hours each wash. The brains were either immediately processed for imaging or stored in DBE at 4°C. The imaging data were collected on the same day.

Brains were imaged using a lightsheet fluorescence microscope (Ultramicroscope, LaVision BioTec, Germany) equipped with an sCMOS camera (Andor Neo, model DC-152Q-C00-FI) and a 2X 0.5 (Olympus MVPLAPO 2XC) objective lens with a 5.6mm working distance of optic correct organic solvent resistance dipping cap. Samples were mounted on the imaging holder, and the merged imaging chamber filled with DBE. Zoom body was adjusted to 1X magnification for imaging. Imaging stacks were acquired using 3 μm steps, and exposure times were varied according to the samples. Each sample was imaged in two different channels, and duplicate images were taken for each channel. The tdT fluorescence was obtained with an excitation filter 560/30nm and an emission filter 625/30nm. Sample autofluorescence was collected for imaging registration with an excitation filter 475/39nm and an emission filter 525/50nm.

#### Whole-brain imaging developmental template transformation and signal detection

To established an age-matched reference template, we first picked one best sample at each age. Since LSFM images were collected in hemispheres, we created synthetic full brains by generating mirror imaged hemispheres and combining them with original hemispheres. We used Elastix^103^ to register all samples in each age to one best represented sample. Then, we used intensity averaging of transformed brains to create a reference template at each age. For anatomical labels, we performed sequential down-registration from P56 Allen Common Coordinate Frameworks to younger ages using Elastix.^51^

Image processing was carried out in Matlab (Mathworks). The background channel of each sample was resized to 20×20×20 μm^3^ stacks. The average autofluorescence of the entire brain in the background channel was determined by masking pixels in the blank space as null values and excluding them from the average. Each full resolution single plane image in both the foreground and background channel was normalized by dividing the intensity of each pixel by the average intensity of background. Both the background and foreground channels were corrected using the flatfield correction algorithm (Matlab) with a 20-pixel area to account for spherical optical distortion introduced by an objective lens. For each full resolution image, the background was subtracted from the foreground to remove vascular and other non-specific signals. A scaling factor was applied to account for overall intensity differences between channels. For samples that were P10 or less, a correction factor of 0.75 was applied to the background image before subtraction. For samples that were P14 to P28, a correction factor of 0.5 was applied to the background image before subtraction. We then calculated the average and standard deviation of the corrected image and used one-half standard deviation above the average value as threshold for signals to generate binarized images. 3D coordinates of detected signal voxels were assigned to the 20×20×20 μm^3^ space by dividing the coordinate values by corresponding resolution changes. Then the accumarray function (Matlab) was used to count the number of signal voxels in 20×20×20 μm^3^ space. The amount of signals was saved as relative percentage occupancy per voxel to represent signal intensity. These values were saved into a 20 μm isotropic voxel image across the brain. To quantify signals in anatomical regions of interest (ROI), we used Elastix to register individual samples to aged matched template brain at 20 μm isotropic resolution.^51,103^ The relative occupancy of summed positive signals per anatomical areas was used to represent signal intensity in each ROI.

All the whole-brain imaging raw and processed data, along with the new developmental brain templates, can be freely accessible at Mendeley data (DOI: 10.17632/m8n8thxxyf.1), BrainImageLibrary (BrainImageLibrary.org).

#### Prenatal Valproic acid (VPA) model NDD model

Timed pregnant mice (E12.5) were IP injected VPA (dissolved in sterile saline) with a dosage of 600mg/kg, and saline was injected as a control.

Self-righting test was performed on VPA and saline treated cohorts using P7 pups as a measure of gross development. Briefly, animals were gently positioned supine with all four limbs pointed upwards. Latency to self-right was measured as the seconds from release until all four limbs touching ground. If the animal had not achieved this position after 60 s, the animal was returned to prone manually, and a time of 60s was recorded. The self-righting test was performed three times with one minute rest in between. The final score is the average latency of the three tests.

Ultrasonic vocalizations emitted by mouse pups can be an early indication of social awareness. On P9, the dam was removed from the housing cage and the pups were transported to a separate location for 30 minutes habituation in the testing room. Animals were then individually placed in a plastic cage, and the cage was placed within a sound-attenuating chamber. Ultrasonic vocalizations were measured for 3 minutes using an UltraSoundGateCM16/CMPA microphone (AviSoft) and SAS Prolab recording software (AviSoft). The number of vocalizations between 33-125 kHz was then counted.

#### Early environmental enrichment rearing

Time pregnant animals (E12.5) were randomly assigned to either standard or early environmental enrichment (EE) housing condition. In the standard housing (SH) condition, one dam and the litter (with a typical litter size around 6) were reared in a standard plastic cage (32.5 × 21 × 18.5 cm) with shredded cotton bedding. The cage was changed weekly. In the EE condition, two pregnant dams and two stranger adult females were housed in a larger plastic cage (50 × 36 × 28 cm), where various sensory stimuli were present, such as variable bedding textures (shredded papers, shredded cotton, corn, shredded wood), visual cues (toys of different colors and shapes), and olfactory cues on wooden cubes. The bedding and environment cues were changed daily. In addition, exploratory behavior was encouraged with the wooden cubes of various sizes, translucent plastic igloos, plastic tubes and tunnels, and a running wheel. The SH- and EE-reared pups were weaned at P28.

For the EE experiments, postnatal SH- and EE-reared mice were monitored between P11 and P15 for eye-opening. For each animal, the opening of one eye was scored as 1, the eye-opening scores ranged from 0 to 2 per animal.

#### Open field

Open field testing was performed on P10 for both EE- and SH-reared mice. Pups and dams were habituated for 30 minutes in the behavior space before testing. During the test, a single pup was placed in the center of an open field chamber and allowed to freely explore for 10 min. The open field chamber was made of polyvinyl chloride with dimensions of 42 x 42 x 42 cm. The chamber was cleaned with 70% ethanol before each test. Pup locomotion and exploration was recorded and analyzed with Any-Maze software. Pup exploration was measured by using a 5×5 (8.4 cm x 8.4 cm) square grid of the chamber floor with the number of individual squares that the pup entered being totaled.

#### Contextual fear conditioning (CFC)

To examine hippocampal-dependent memory ability, CFC behavior testing was conducted on P17, P20, P24, and P28, with all training taking place before weaning. The animals and dam were habituated to handling in the pre-testing area for 3 days prior to CFC training. The CFC training and recall paradigms took 5 minutes and occurred in a testing room adjacent to the habituation area. Behavior tastings were performed at a consistent time of day. During training, animals were placed in a custom context box and allowed to explore for 1 min 58 seconds before receiving the first of a series of three 0.5 mA shocks, each lasting for 2 s with 1-minute intervals between shocks. Animals were removed from the context box one minute after the final shock and returned to the home cage with the dam.

The CFC recall tests occurred 1 day and 7 days after training. All animals were only trained once and tested twice. The following training/1 day recall/7 day recall groups were used for data collection: P17/P18/P24, P20/P21/P27, P24/P25/P31, and P28/P29/P35. CFC recall involved the animals and dam being returned to habituation area and recall testing occurred individually. Animals were placed in the same context as training for 5 minutes without receiving any shocks. Contextual memory was assessed by measuring the freezing behavior displayed in the animals within this context. Freezing was defined as the total absence of movement apart from respiration. Video recordings were also made to confirm accuracy behavior quantification. All the recordings and analysis were performed blinded.

#### Sequencing data analysis

##### scRNA-seq data processing, clustering, and data integration

Cellranger was used to demultiplex Libraries BCL files, analyze feature barcodes, and annotate resulting transcripts to the mm10 reference genome (UCSC genome browser), with modification to contain the tdT sequence at the Rosa26 loci.^32^ We identified possible doublets using Scrublet on each library individually and then concatenated all 4 libraries in scanpy.^92,93^

Quality control thresholds were determined using scanpy. Cells were excluded if they were predicted to be a doublet or had a doublet score of >0.35. We excluded cells with greater than 60,000 reads (UMI counts) and less than 2,500 genes. All subsequent analysis was performed in R (v4.1.0).

Seurat V4 was then used to determine cell types.^94^ SCTransform was first implemented to scale gene expression and in this step variance due to mitochondrial fraction was regressed out.^104^ Variable features were selected to create PCs using VST method of FindVariableFeatures. 30 PCs were used to find neighbors, clusters, and UMAP dimensions. Clusters were then classified into cell types based on the expression of the indicated genes (Figure S5C).

Next, we took steps to limit the potential for downstream analysis to be biased by over-clustered data. To do this we first split the single-cell dataset into neurons and non-neurons and generated new PCAs for the neuron datasets to better capture the genetic diversity of this important cell type. For both datasets, we used the FindTransferAnchors and TransferData functions to project cell cluster identities back onto the same single-cells (using the same data as both reference and query sets). We found that the projected cell types were very consistent with the originals (neurons: 95.0% accuracy, non-neurons: 99.4% accuracy). As desired, some cell clusters were lost when projecting, indicating these groupings were likely the result of over-clustering (Figure S5B). A few remaining clusters were removed due to poor data quality: olig neuron, exclude (microglia), and high-mt GCs. Moving forward the projected cell clusters were used as the default (Figure S5C).

With the filtered cell population we calculated new PCs and UMAP using 20 PCs. This data we evaluated using the standard quality controls of mitochondrial fraction, number of genes, and number of counts (Figure S5A). We used the SCTransformed tdT counts to binarize tdT status at a threshold of 2 (Figure S5D).

Finally, the granule cells (GCs) were isolated based on cell cluster and a few straggler cells were eliminated based on UMAP filters (5,562 GCs). We then normalized the SCT counts with edgeR to calculate CPM and applied a log2 transformation.^95^ This data was then reformatted as a SingleCellExperiment to be compatible with downstream analysis.^105^

We next filtered out genes that were present in <25% of GCs, leaving 7,904 genes (Table S5). These are the genes and expression values used for differential expression analyses.

##### Cluster marker gene identification

We ran Seurat’s FindMarkers function on SCTransformed RNA values to identify cluster-specific gene markers. Results were filtered to genes that were present in at least 25% of all GCs then filtered by adjusted p value < 0.0001 and average log2 fold change >0.5. This produced 484 nascent GC markers, 14 transition GC markers, 102 growth GC markers, and 117 synaptic GC markers. The top markers for each group were selected based on the following criteria: 10 genes with the lowest adjusted p value, 10 genes with the lowest minimum adjusted p value between SH and EE, 10 genes with the highest average log2FC, and 10 genes with the greatest difference in percent expression (vs other GC clusters). This produced 94 nascent GC markers, 13 transition GC markers, 25 growth GC markers, and 15 synaptic GC markers. Due to the large number of nascent GC markers with an adjusted p value equal to 0, nascent GC markers were re-generated without adjusted p value filters. This produced a total of 65 unique genes that are the top markers differentiating GC clusters (Figure S5G).

##### Trajectory analysis

To reconstruct the cell-stage transition during GC development, we used the Slingshot package to generate pseudotime estimates from gene expression.^96^ Pseudotime analysis on PC representations of log2(CPM) gene expression^97^ indicated a single pseudotime trajectory that we used as an estimate of developmental maturation (Figure 6d). When examining pseudotime along with other experimental outcomes, we used a binned version of the pseudotime measurement. For statistical comparison of tdT pseudotime at different ages and visualization as a cumulative distribution plot, 20% of the GCs were randomly sampled (P14 n= 223 (29 tdT+), P20 n= 405 (141 tdT+), P24 n= 479 (246 tdT+). Kolmogorov-Smirnov tests were performed with a two-sided alternative hypothesis.

##### Differential gene expression

We examined the CPM normalized log2 counts of 7,904 genes for significant differences based on our experimental variable of binarized tdT status. The MAST package was used to implement zero-inflated linear models testing for the experimental condition with consideration of library depth as a covariate.^98^ For all differential expression analyses, significance was evaluated at an FDR < 0.0001. To consider the interaction of our experimental variables of tdT status with the biological variables of age and GC group, we split the GC population by these variables and created a separate model for each level. For age-specific analysis significance further required an absolute log2FC of 0.3. For cluster-specific analysis significance further required an absolute log2FC of 0.1. The UpSetR package was used to visualize DEGs across ages.^99^

##### Gene set enrichment analysis (GSEA)

The clusterProfiler package was used for GSEA of gene ontology biological processes terms.^100^ For all GSEA performed, our own whole-DG dataset was used to generate the universe background (19,025 genes). For all GSEA performed, only results with adjusted p<0.05 were considered.

Genes used for cluster-specific GSEA were derived from cluster markers (102 growth GC markers, 117 synaptic GC markers). Given the inflated number of unique nascent GC markers, the 134 top markers were used. Transition GCs were not included in GSEA as this cluster did not demonstrate a sufficiently unique transcriptome from the other clusters. These gene lists generated 29 nascent GC terms, 32 growth GC terms, and 44 synaptic GC terms. The top 10 non-redundant terms were chosen from each analysis and merged using clusterProfiler. These results are summarized in Figure 6F, where the color of the node indicates which analysis produced that significant term. Nodes with split colors indicate the presence of the term in more than one analysis and the proportion of the colors within the node denotes the proportion of genes contributed by each analysis.

For tdT GSEA, we were more interested in a thorough characterization of enriched biological processes and so fewer filters were applied to the resulting terms. GSEA for tdT-labeled cells was performed on 245 genes increased in tdT-labeled cells at all P14, P20, and P24.

##### Activity-regulated genes (ARGs)

We compiled a list of ARGs based on three published datasets, indicating activity-induced transcription based on prolonged *in situ* stimulation,^87^ visual experience^86^ and a dependence on fos transcriptional activity.^106^ We isolated 423 genes upregulated based on sensory experience, 168 genes upregulated based on *in situ* stimulation, and 123 genes that were increased in more than one experimental manipulation in Yap 2021. Of those, the following gene sets were also expressed in more than 25% of our GC samples: 219 sensory experience ARGs, 58 *in situ* ARGs, and 83 fos-associated ARGs.

We further limited this list to 21 depolarization induced genes (experience-dependent and *in situ*, with 19 also showing fos association), 64 fos-associated experienced-dependent ARGs, and 22 fos-associated stimulation-induced ARGs. Scater was used to generate the heatmap of these 107 ARG genes across pseudotime.^97^

## Notes

### Competing Interest Statement

The authors have declared no competing interest.

